# Genetic variation exists within *Zea mays* to influence unsustainable nitrogen cycling microbiome function

**DOI:** 10.1101/2023.09.19.558450

**Authors:** Alonso Favela, Martin O. Bohn, Angela D. Kent

**Affiliations:** Program in Ecology, Evolution, and Conservation Biology, University of Illinois at Urbana-Champaign, Urbana, IL 61801; Department of Ecology and Evolutionary Biology, University of California Irvine, Irvine, CA, 92697, USA; Department of Crop Sciences, University of Illinois at Urbana-Champaign, Urbana, IL 61801; Department of Natural Resources and Environmental Sciences, University of Illinois at Urbana-Champaign, Urbana, IL 61801

**Keywords:** Microbiome, Rhizosphere, Maize, Selection, Nitrogen Cycle, Agro-ecosystem, Sustainability, Nitrification, Denitrification

## Abstract

Overuse of synthetic nitrogen fertilizers in agroecosystems causes environmental pollution and human harm at a global level. Nitrogenous fertilizers provide a short-lived benefit to crops in the agroecosystem, but stimulate microbially-mediated nitrification and denitrification, processes that result in N pollution, greenhouse gas (GHG) production, and reduced soil fertility. Recent advances in plant microbiome science suggest that plants can modulate the composition and activity of rhizosphere microbial communities. These rhizosphere communities act as an extended phenotype, primed by genetic variation in the plant host. Genetic variation in traits (e.g., plant secondary metabolites, root architecture, immune system, etc.) act as mechanistic selective agents on the composition of the microbiome. Here we attempted to determine whether genetic variation exists in *Zea mays* for the ability to influence the extended phenotype of rhizosphere soil microbiome composition and function. Specifically, we determined whether plants’ influence on soil nitrogen cycling activities was altered by plant genetics and thereby allowing it to be incorporated into breeding practices. To capture an extensive amount of genetic diversity within maize we sampled the rhizosphere microbiome of a germplasm chronosequence that included ex-PVP inbreds, hybrids, and teosinte (*Z. mays* ssp*. mexicana* and *Z. mays* ssp. *parviglumis*). We observed that potential N cycling processes were influenced by plant genetics. Teosinte and some hybrid genotypes supported microbial communities with lower potential nitrification and potential denitrification activity in the rhizosphere, while inbreds stimulated/did not inhibit these undesirable N-cycling activities. These potential differences translated to functional differences in N_2_O production, with teosinte plots producing less GHG than maize plots. Furthermore, within these *Zea* cultivars we found that plant genetics explained a significant amount of variation in the microbiome, particularly among different nitrification and denitrification functional genes within the community. We found that potential nitrification, potential incomplete denitrification, and overall denitrification rates, but not abundance of N-cycling genes of rhizosphere soils were influenced by growth stage and plant genetics. Taken together, these results suggest that crop selection can lead to changes in root phenotypes that could suppress unsustainable N-cycling processes. Reintroducing stress-adapted and “wild” root characteristics into modern germplasm may be a way to manipulate soil microbiomes at both a composition and functional level to improve sustainability.

## Introduction

More than half the world’s population depends on crops grown with synthetic nitrogen (N) fertilizers (Bowles et al., 2018). Regrettably, most of these synthetic N fertilizer inputs escape the agroecosystem, degrade natural areas, and harm human health (Vitousek et al., 2013; Zhang et al., 2015). Stimulation of soil nitrogen cycling microorganisms (i.e. nitrifiers and denitrifiers) is a major contribution to “leaky” agricultural systems (Kuypers et al., 2018). To improve the sustainability of agricultural systems, we need to understand how ecological drivers, such as plant-microbe interactions, influence soil’s nitrogen cycling microorganisms and the movement of nitrogen, and how this can be managed to reduce nutrient losses (Moreau et al., 2015, 2019).

Genetic variation within crop species has been shown to play a significant role in plant-microbiome assembly and recruitment (Peiffer et al., 2013a; Bouffaud et al., 2016; Walters et al., 2018). Across large-scale and multi-year field trials, researchers find consistent sets of heritable core microbial taxa associated with specific plant genotypes (Walters et al., 2018; Xu et al., 2018). These taxonomic assembly differences are functionally relevant, as microorganisms contain a diverse biochemical repertoire that allows plants to escape nutrient, drought, and pathogen stress (Philippot et al. 2013; Compant et al. 2019; Trivedi et al. 2020). In the rhizosphere, plants exude chemical cocktails of metabolites, the production of which is directed by the plant genome. These exudate traits, in tandem with root phenology and physiology, act as an ecological filter that has direct fitness consequences for the surrounding soil microbial communities (Canarini et al., 2019; Huang et al., 2019). Soil microorganisms that are phytochemically competent to the ecological filter of the rhizosphere survive and persist near the plant root (Philippot et al. 2013). It follows, then, that the ecological filtering for rhizosphere occupancy that arises from genotypic variation directing microbiome selection will have direct consequences on biogeochemical cycling activities carried out by the rhizosphere microbiome.

Rhizosphere soil microorganisms are key contributors to essential ecosystem functions. Processes such as nitrification and denitrification are primarily controlled by microorganisms in the soil and can result in a considerable loss of nitrogen from an ecosystem (Philippot et al., 2007; Davidson et al., 2012). Microorganisms and their suite of functions are extremely diverse, and variation in assemblages of species likely plays a critical role in shaping how ecosystem processes occur in ecological settings. In many ways, soil microorganism biodiversity is a foundational gatekeeper of the pathways by which nutrients can enter and exit an ecosystem. Furthermore, emerging research is beginning to show that genetic variation within plant species can have a considerable role in driving both soil community composition and biogeochemically-relevant microorganisms (Subbarao et al. 2013; Pérez-Izquierdo et al. 2019). For example, we have shown that historic selection on plant genotype can drive the assemblage of the nitrogen cycling rhizosphere microbiome, both by changing the taxonomic composition and representation of nitrogen-cycling functional genes (Bouffaud et al., 2016; Favela et al., 2021). Yet we and others have lacked evidence linking microbiome changes to altered nitrogen cycling processes within the ecosystem. Here, we attempted to address this in an agroecological field setting. Specifically, we sought to determine whether plant genetic variation involved in root phenotypes is an important explanatory variable for understanding changes in the soil microbiome, particularly microbial functional groups, and to connect variation in this extended phenotype back to changes in important nitrogen cycling processes.

To evaluate the effect of plant genetic variation across root phenotypes influence on nitrogen cycling microbial functional groups, we grew a diverse panel of *Zea mays* (including elite inbreds, their hybrids, and wild teosintes) and measured their contribution to differences in microbial community assembly and nitrogen cycling processes. By doing this in a highly replicated manner within a single field, we were able to control for stochastic edaphic factors and parse the plant genetic contribution driving soil microbiome function in an agronomically relevant field setting. Previously, our research in elite inbred maize suggested that breeding of maize resulted in a narrowing and loss of sustainable N-cycling microbiome functions through the 20^th^ century (Schmidt et al., 2019; Favela et al., 2021). Informed by this prior research, we included maize hybrids to estimate if microbiome functions would be regained through heterosis (Wagner et al., 2021). In addition to this, within *Zea mays*, we found microbiome recruitment and N-cycling functional groups differed the most between modern inbred maize and wild teosinte (Favela et al., 2022). Therefore, wild *Zea* was included as an outgroup to evaluate the influence of pre-domestication genetics and their ecological filtering traits on microbiome assembly and function (Brisson et al., 2019; Schmidt et al., 2020). These treatments give us an understanding of how much plant genetic variation is necessary to induce changes in the microbiome of soils and provide insight into how domestication and breeding for performance in high N environments altered microbiome functions. With this information, we hoped to better understand how genetic alterations in maize can have cascading effects in plant microbiome recruitment and nitrogen cycling activity. Understanding plant genetic contributions to these functional processes in an agroecological field setting is critical for improving the sustainability of maize production.

## Methods

### Field Design

Field plots were located at the Crop Sciences Research and Education Center (CSREC) - South Farms at the University of Illinois, Urbana-Champaign, IL (40°03’30.4”N 88°13’50.4”W). We used a panel of *Zea* cultivars that encompassed modern ex-PVP inbreds, their hybrids, and the wild progenitor of maize, teosinte (27 genotypes in total: Description of genotypes in Table **S1**). Teosinte was represented by the subspecies *Zea mays* spp. *mexicana* and *Zea mays* spp*. parviglumis*, with three genotypes per subspecies). Across our field, each cultivar was replicated four times. The replications were arranged in a randomized complete block design (Fig. **S1-2**). Each maize plot contained four rows with 15 plants per row. Maize inbreds and hybrids were machine planted on 4/30/2017 while teosinte cultivars were hand planted 5/19/2017. Due to seed limitations teosinte was planted in 2-row plots. Fields are managed by the CSREC in a Corn-Soy rotation, tilled yearly, and plant density of 32,000 plants/acre. Fertilization application was applied homogenously across all plots and match those on a typical conventional agricultural farm in Illinois (82 N kg/acre, 35 P kg/acre, 23 K kg/acre).

### Sample collection

Plots were sampled three times. Plants were sampled in approximately in the V4 (Maize and teosinte: 6/5/17), V6 (Maize: 6/20/17, Maize and teosinte:7/20/17), and R2 (Maize:7/12/17, Teosinte: 8/11/17) growth stages across the growing season. Staggered sampling of teosinte and maize at later timepoints was done to control for differential growth rates among these genotypes. This sampling disconnect was accounted for in statistical models for nitrification and denitrification by modeling growth stage as a fixed factor and sampling date as a random factor. We found minor influence of both. At each sampling event, four individual plants per plot were sampled and combined into a composite sample. Individual plants were not resampled, thereby maintaining independence of sampling timepoints. To account for the size of the experiment, true “rhizospheres” were not collected. Instead, root zone soil was collected as a proxy of the rhizosphere. Root zones soils here consisting of a mixture of rhizosphere and bulk soil present in the immediate proximity of the plant. This method allowed us to sample enough soil for nitrogen cycling assays and molecular work without having to carry out time consuming rhizosphere soil extractions from the roots. Samples consisted of a soil core (10 cm depth) obtained from the root zone of the plant (2 cm away from base of stem). Composite samples were placed on ice until they were transported to the lab. Processing of soil cores before assays and molecular work consisted of removal of all root tissue present in sample and homogenization of soil cores. Root tissue present in root zone soils were removed before further processing. Once in the lab, soils were refrigerated at 4°C awaiting potential nitrification and potential denitrification assays (within 5 hours). Aliquots for DNA extraction were frozen immediately. The frozen DNA aliquots were placed into 15 mL centrifuge tubes and lyophilized before DNA extraction (0.5g total) using the FastDNA for Soil DNA extraction kit (MPBio, Solon, OH). Soil samples were collected at the end of the season (9/15/17) for nutrient analysis carried out by Waypoint Analytical (Champaign, IL, USA).

### Potential nitrification assay

The potential nitrification assay was developed and modified from (Schinner et al., 1996). This assay was performed at substrate saturation and values presented should be interpreted as the maximum potential rate of transformation of ammonium to nitrite, the first-rate limiting step of nitrification. In principle, this assay uses ammonium sulfate as the substrate for the first step of nitrification during a 5-hour incubation. Nitrite products released during the incubation period were extracted with potassium chloride and concentration is determined colorimetrically at 520nm. Sodium chlorate was added to the assay to inhibit nitrate oxidation during the incubation period. Sample tubes were incubated on a rotatory shaker for 5h and control tubes were stored at -20°C for 5h. After incubation and thawing, KCl was used to extract nitrite from both samples and controls. Potential nitrification rates were arithmetically adjusted by initial soil moisture, soil weight, % dry matter, and initial nitrite in the sample. Potential nitrification data is presented as (log (ng N g^-1^ hr^-1^)) and percent change ((population mean – genotype mean) divided by population mean).

### Potential denitrification rates by acetylene-inhibition assay

Potential denitrification assays were carried out using a modified version of previously described assays (Schinner et al., 1996; Peralta et al., 2016). Field-moist root-zone soil samples were incubated under anaerobic conditions in the presence of acetylene for 3h at 25°C. The assay was performed on 25g of root-zone soil in 125-ml glass Wheaton bottles. Incubations were carried out at substrate saturation of carbon (dextrose) and nitrogen (nitrate). Chloramphenicol (10 mg/L) was added to the incubation to act as a bacteriostatic agent to prevent further microbial growth and protein synthesis. The incubation bottles were purged of oxygen with either helium or acetylene. Helium samples were used to estimate the amount of incomplete denitrification (N_2_O) produced during the assay. Acetylene purged samples were used to measure overall denitrification (N_2_O + N_2_). Acetylene is a commonly known inhibitor of nitrous oxide reduction. Initial and final gas samples were collected at the start and end of the incubation period. Initial and final nitrous oxide in gas samples was quantified using a GC-2014 Gas Chromatograph (Shimadzu, Kyoto, Japan) with an electron capture detector (GC-ECD). Potential denitrification rates were arithmetically adjusted by initial soil moisture, soil weight, % dry matter, sample volume, and headspace. Potential denitrification data is presented as (log (ng N g^-1^ hr^-1^)) and percent change ((population mean – genotype mean) divided by population mean)).

#### Carbon Substrate Utilization

Carbon substrate utilization assay was carried out using Biolog EcoPlates (Biolog Inc., Hayward, CA, USA). Biolog EcoPlates are a simple method to characterize the metabolic functions of microbial populations. Plates contain 31 different carbon substrates that can be used as the sole source of carbon. Each substrate is bound to a tetrazolium dye that changes colors once carbon compound is degraded. Assays were carried out with soils collected from the 7/20/17 sampling timepoint (represented by the R2 timepoint). Root zone soils (0.5g) were diluted (1:4) in PBS, vortexed and centrifuged. Soil mixture supernatant (600 uL) was further diluted (1:25) in PBS. The diluted soil mixture was then added to the microplates and incubated for 5 days at room temperature. Absorbance of substrates was measured every 24 hours using an Epoch microplate spectrophotometer (Santa Clara, CA, USA). Microbial metabolism was calculated as suggested in (Classen et al., 2003). This comparison focused on inbred B73, hybrid check1, and PI566677 teosinte (4 replicates per genotype). Data used in analysis consisted of carbon substrate usage (average well development) across 4 technical replicates at end of 5 day incubation. The SIMPER procedure in the ‘vegan’ R package was used to determine differences in substrate utilization across treatments (Oksanen, 2017).

### Microbial Community Amplicon Sequencing

For this experiment, we characterized the microbiome and diagnostic functional genes related to transformations that occur in the nitrogen cycle: nitrification, and denitrification. Amplicon sequencing was performed on bacterial and archaeal 16S rRNA genes, fungal ITS2, *amoA, nirS, nirK,* and *nosZ* genes. The Fluidigm Access Array IFC system was used to prepare sequencing amplicons. This method allows for the simultaneous amplification of target functional genes using multiple primer sets (Fluidigm, San Francisco, CA). DNA sequencing was performed for bacterial, archaeal, and fungal amplicons using an Illumina NovaSeq Sp flowcell with 2 x 250 bp reads (Illumina, San Diego, CA). Primer information is provided in supplemental Table **S2**. Fluidigm amplification and Illumina sequencing were conducted at the Roy J. Carver Biotechnology Center, University of Illinois (Urbana, IL, USA). Fast Length Adjustment of Short reads (FLASH) (Mag and Salzberg, 2011) software was used to merge paired-end sequences from bacterial and archaeal 16S rRNA genes. Due to the amplicon size for some functional genes, only forward read sequences were used. Once FLASH merging was performed, files were filtered by quality using the FASTX-Toolkit (Hannon, 2014). Reads that did not have a minimum quality score of 30 across 90% of the bases were removed. Using the FASTX-Toolkit, *nirK* sequences were trimmed to the amplicon size of 165-bp.

Once quality preprocessing was performed, FASTQ reads were converted to FASTA format. Using USEARCH-UPARSE version 8.1 (Edgar, 2010), sequences were binned into discrete OTUs based on 97% similarity and singleton DNA sequences were removed. Quantitative Insights into Microbial Ecology (QIIME) was used to generate OTU tables for downstream statistical analysis and to assign taxonomic information, this is done with a combination of the UCLUST algorithm and SILVA 138.1 database (DeSantis et al., 2006; Caporaso et al., 2010; Edgar, 2010). Once taxonomy was assigned, chloroplast and mitochondrial OTUs were removed from the dataset. Rarefaction was performed to correct for differential sequencing depth across samples. Functional gene sequences were also assigned using QIIME (Caporaso et al., 2010) with the BLAST algorithm (Altschul et al., 1997) and custom gene-specific databases generated from reference sequences obtained from the FunGene repository (http://fungene.cme.msu.edu/) (Fish et al., 2013). All OTU tables used in statistical analyses were generated in QIIME. Singleton OTUs were filtered prior to statistical analysis.

The number of raw reads generated from sequencing run, reads present after quality filter, and the rarefaction level are reported in supplemental Table **S3**. Amplicon sequence data for 16S rRNA genes, fungal ITS2 region, and N-cycling functional genes is available for download on the NCBI SRA database at accession number: PRJNA789877. (https://www.ncbi.nlm.nih.gov/bioproject/PRJNA789877/). Code for sequence processing and statistical analysis is available in GitHub (https://github.com/favela3/Maize.N-cycle.Function).

### Statistical Analysis

Statistical analysis was performed in R, with the packages ‘Vegan’, ‘ASReml’, and ‘WGCNA’. ‘Vegan’ was used to perform multivariate statistical comparisons for microbiome data among experimental treatments (Oksanen et al., 2007; Langfelder and Horvath, 2008; Butler et al., 2017). ‘ASReml’ was used to perform univariate comparisons among genotypes and cultivar classes potential nitrification, potential denitrification and nitrogen cycling qPCR. WGCNA was carried out to compare multivariate microbiome data to univariate nitrogen cycling function data. Model factors used in statistical analysis were growth stage, sampling date, the location of the block, the row of block position, range of block position, the genotype within the block, and the interaction between genotype and time (combined growth stage and sample date). A typical model of analysis is displayed below:

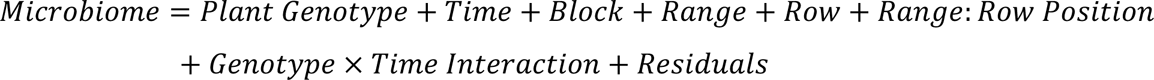

In PERMANOVA models, block factor was constrained in permutations. For the PERMANOVA models, sampling date and growth stage are combined as a time factor. In the ASReml mixed effect models for potential nitrification and denitrification, plant genotype and the genotype × growth stage interaction were treated as fixed factors, while all other factors (block, range, row, sampling date) were treated as random factors.

### Quantifying nitrogen cycling functional groups

Quantitative PCR (qPCR) was carried out to determine the abundance of functional genes in each of the root zone microbial communities. Specific target amplification (STA), explained in Ishii et al. (2014), was carried out on samples and standards to increase template DNA for amplification. STA and qPCR master mix recipes from (Edwards et al., 2018) were used for all samples. STA product and qPCR master mix were loaded into the Dynamic Array™ Microfluidics Fluidigm Gene Expression chip, where amplification and quantification of functional genes were carried out simultaneously (Fluidigm, San Francisco, CA). All samples and standards were analyzed in 12 technical replicates. Fluidigm Real-Time PCR Analysis software version 4.1.3 was used to calculate gene threshold cycles (C_T_). C_T_ values were converted to gene copy number using gene length and standard curves. All Fluidigm qPCR was conducted at the Roy J. Carver Biotechnology Center (Urbana, IL, USA). The final copy number of each functional gene amplicon was standardized by the ng of template DNA in the qPCR reaction.

### In situ N_2_O flux measurements

Net soil-atmosphere N_2_O fluxes were measured weekly from 6/20/17 to 8/23/17, samples were collected for a total of 6 weeks. As gas flux measurements are laborious and time consuming, sampling was targeted during plant peak primary plant growth and focused on the plant treatments that were hypothesized to have the largest effect on the microbiome function based on previous studies (Favela et al. 2021b). Specifically, the comparison focuses on B73 inbred, check1 hybrid, and PI566677 teosinte. Flux measurements were measured using static flux chambers as described in USDA-ARS GRACEnet Project protocol (Parkin and Venterea, 2010). Chambers were installed in the field during the first sampling timepoint and remained in place throughout the maize growing season. Chambers consisted of two-pieces: PVC pipe with a 30 cm diameter (base installed 20 cm into soil), and sampling lids (10 cm in height). Gas sampling events occurred in the mornings between 10 am-noon; during this time 15 mL of gas were collected from chambers every 10 minutes for 30 mins. Samples were stored in evacuated aluminum crimp-top glass vials with a chlorobutyl stopper and sealed with clear silicone to prevent sample leakage. Gas samples were later quantified using a GC-2014 Gas Chromatograph with an electron capture detector (GC-ECD) (Shimadzu, Kyoto, Japan). Standard curves were used to quantify the amount of N_2_O in gas sample. N_2_O samples were corrected using ambient temperature and moisture conditions of the collection day. Four sampling timepoints were used to determine the rate of N_2_O flux (mg m^-2^ min^-1^). While linear interpolation was used to estimate cumulative N2O flux (mg L^-1^) over the growing season (Parkin, 2008).

## Results

### Nitrogen Cycling Genes Community Across the Agroecosystem

From our analysis of nitrogen cycling functional genes, we observed 210 archaeal *amoA* OTUs, 98 bacterial *amoA* OTUs, 21022 *nirK* OTUs, 2607 *nirS* OTUs, and 7294 *nosZ* OTUs (DNA sequencing quality is described in Table **S3**). In response to genotype, the overall microbiome and 4 of 5 nitrogen cycling genes showed statistically significant changes in community membership (Fig. **1**; Table **S4**), 1 of 5 nitrogen cycling genes changed in abundance (Table **S4.1**). Conversely, plant classification (i.e., inbred, hybrid, teosinte) affected the composition of 4 of 5 nitrogen cycling genes, and the abundance of 2 of 5 nitrogen cycling genes (Table **S4.2**). Additionally, plant genotype and genotype classification interactions with time had a significant effect on N-cycling composition and functional gene abundance (Table **S4.1-2**; Fig. **S5-6**).

**Figure 1.**
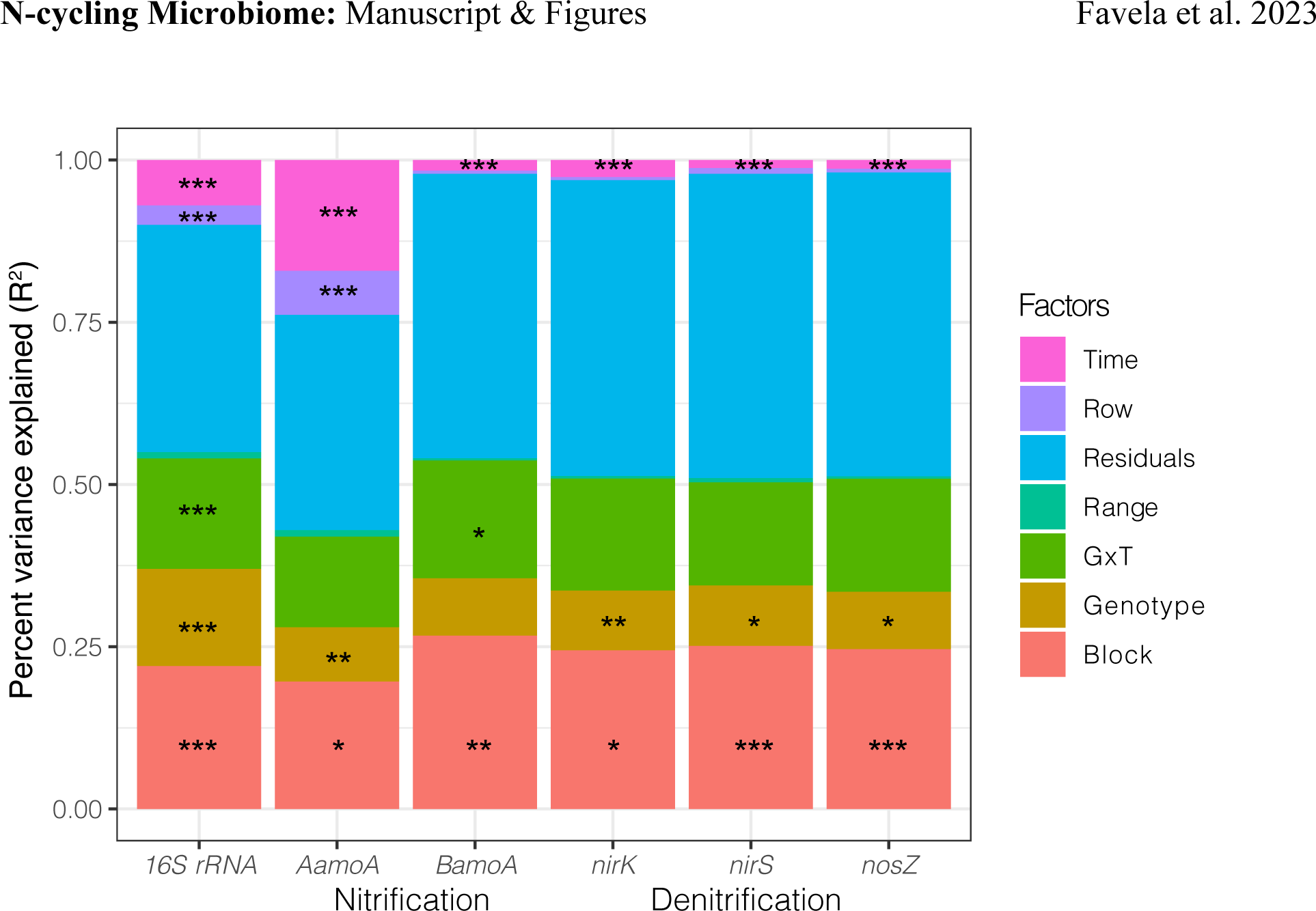
PERMANOVA results for the overall microbiome (16S rRNA) and different nitrogen cycling functional genes included in this study (nitrification: *AamoA* – archaeal *amoA* [ammonia monooxygenase], *BamoA* – bacterial *amoA*; denitrification: *nirS* and *nirK –* nitrite reductase, *nosZ –* nitrous oxide reductase). The y-axis shows R^2^, percent variance explained by the treatment factor, and the x-axis shows the functional genes tested. (* – P<0.05, ** – P<0.01, *** – P<0.001).

### Nitrification genes and potential function

Recruitment of nitrifiers (indicated by gene sequences for bacterial and archaeal ammonia monooxygenase – *amoA*) was not significantly impacted by plant genotype and plant classification. While plant genotype explained a small but significant amount of variation for archaeal *amoA* (R^2^=0.08, p<0.001, Fig. **1**, Tables **S5.1-3**), there was not a significant change in community composition of bacterial ammonia oxidizers in response to genotype (p=0.16, Table **S4, S5.4-6**). Regarding abundance, neither archaeal nor bacterial ammonia oxidizers were significantly influenced by plant genotype (archaeal *amoA* p=0.61, bacterial *amoA* p=0.99, Table **S6.1-7**). Plant classification showed the same patterns as genotypes (Fig. **S5,** Table **S4, S5, S6**). Potential nitrification rate (log (ng N g^-1^ hr^-1^)) of root zone soils was influenced by both time of growth stage, genotype, and plant classification (Fig. **2a, S3a**, Table **S7.1-2**). Specifically, teosinte genotypes had lower potential nitrification rate by 9% compared to inbred maize which, on average, stimulated potential nitrification rates by 4% (means difference of 13%, Fig. **3a, S9.1**). It should also be noted that a considerable amount of variation in potential nitrification rates could be attributed to plant genotype (p<0.05, Table **S7**).

**Figure 2.**
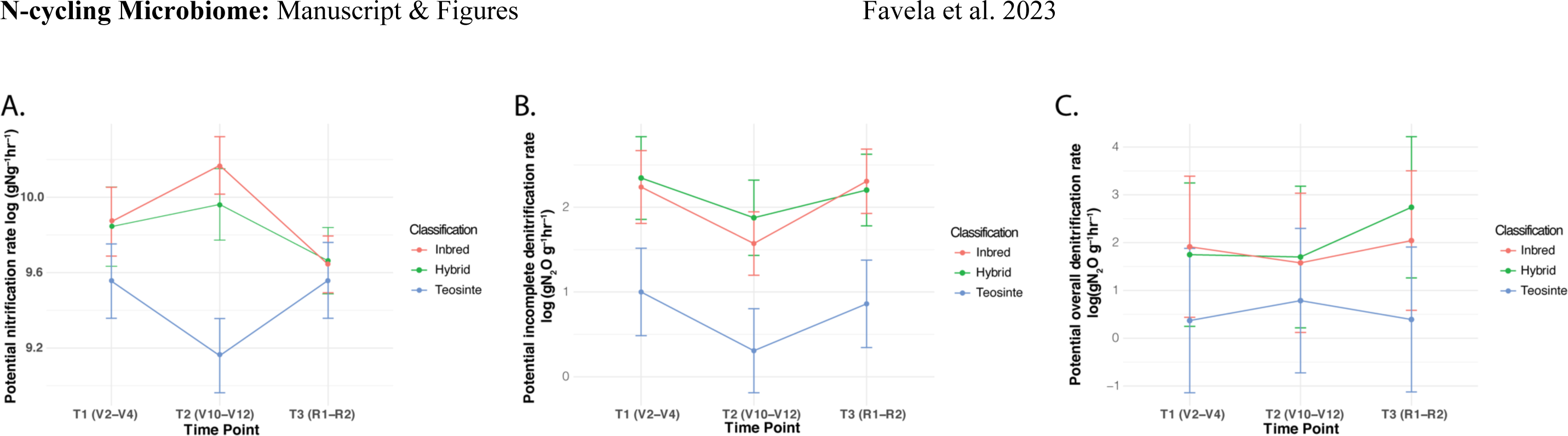
Seasonal variation in potential nitrification and denitrification rate compared among germplasm group across the season. LS Means and Standard Error were calculated using ASReml-R. **A.** Potential nitrification rate (log (ng N g^-1^ hr^-1^)) across the three sampling time points averaged over plant classification. **B.** Potential incomplete dentification (log (ng N_2_O g^-1^ hr^-1^)) across the growing season averaged over plant classification, no differences in among plant classifications was observed, but potential denitrification rates increased slightly across the season. **C.** Potential overall dentification (log (ng N_2_O g^-1^ hr^-1^)) across the growing season averaged over plant classification, no differences in among plant classifications was observed, but potential denitrification rates increased slightly across the season. Statistical tests associated with figures are presented in supplemental materials Tables S7-8.

**Figure 3.**
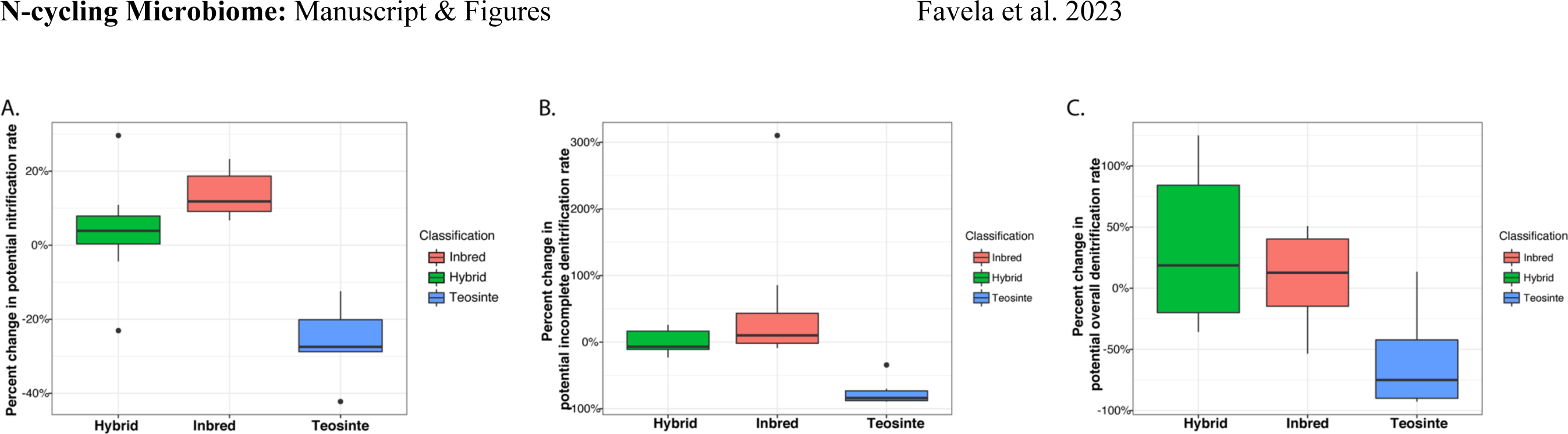
Plant classification influences rhizosphere N-cycling activities. **A.** Average genotypic effect across all timepoints of hybrid, inbred, and teosintes genotypes on the potential nitrification (log (ng N g^-1^ hr^-1^)) determined across the population. Statistical analysis for both figures can be found in supplemental tables S7-9. **B.** Average genotypic effect across all timepoints of hybrid, inbred, and teosintes genotypes on the log of potential overall denitrification (log (ng N_2_O g^-1^ hr^-1^)) determined across the population. Statistical analysis for both figures is included in supplemental tables S7-9. **C.** Average genotypic effect across all timepoints of hybrid, inbred, and teosintes genotypes on the log of potential incomplete denitrification (log (ng N_2_O g^-1^ hr^-1^)) determined across the population. Percent change calculation described in Methods. Statistical analysis for both figures is included in supplemental table S7-9.

### Denitrification genes and potential function

All the denitrification genes surveyed were significantly different among genotypes and plant classification (Fig. **1, S6**, Tables **S4, S5.7-15**). Communities of denitrifiers possessing the cytochrome cd_1_-type nitrite reductase (encoded by *nirS*) or the copper containing nitrite reductase (encoded by *nirK)* both varied significantly among plant genotypes (*nirS:* R^2^=0.09, Fig. **1**, Table **S5.8**; *nirK:* R^2^=0.09, p=0.003, Fig. **1**, Table **S5.11**) In addition, *nosZ,* the gene that encodes typical nitrous oxide reductase, crucial for the consumption of N_2_O, was found to be affected by plant genotype (R^2^=0.09, p=0.011, Fig. **1**, Table **S5.14**). Quantitative PCR of denitrification genes showed no difference in the abundance of genes in the root zone across plant genotype and largely for plant classification (Fig. **S5-6**, Table **S4**). One exception to this was *nosZ,* which was observed to be altered by plant classification (p<0.05, Table **S5.13**, Fig. **S4e**). Similar to the observations for bacterial *amoA*, the denitrification genes also showed a strong and significant interaction between plant classification and time (Fig. **S5-6**). During the first sampling time point (V2-V4), the teosinte root zone microbiome contained similar denitrification gene abundance, had greater levels of dentification genes during the final (R1-R3) sample point, and finally, had lower numbers of denitrification genes compared to inbred and hybrid maize by the end of season. Full analysis of plant classification effects and additional genotype models on denitrification genes is presented in Supplemental Tables S5-6. Conversely, we found variation in the potential incomplete denitrification (N_2_O) and overall denitrification (N_2_O + N_2_) rates (log (ng N g^-1^ hr^-1^)) of root zone soils to be consistently influenced by genotype and plant classification, but not growth stage (Fig. **2b, S3b**, incomplete: p<0.001, overall: p<0.001, Table **S8.1-4**). For potential incomplete denitrification (log (ng N g^-1^ hr^-1^)), teosinte genotypes had lower activity by 75% and inbred genotypes on average stimulated potential complete denitrification by 32% (mean difference 102%, Fig. **3b**, Table **S9.3**). On average, teosinte genotypes had lower the log of overall denitrification by 59% compared to inbred maize, which stimulated it by 4% (mean difference 63%, Fig. **3c**, Table **S9.2**).

### Static N_2_O flux Chambers

To estimate whether our potential denitrification and nitrification rates were reflected in ecosystem flux differences, we placed static flux chambers in blocks with three of our genotypes inbred, hybrid, and teosinte. From these static chambers, we found that over the growing season, the root zone soil of the teosinte genotypes produced significantly less cumulative N_2_O production (mg L^-1^) (t=2.01, df =29, p=0.05, Fig. **4a**) and had lower N_2_O flux (mg m^-2^ min^-1^) (t=2.09, df=33, p=0.04) compared to the inbred genotype. Hybrid plots were not significantly different in N_2_O production from inbred and teosinte (B73: t=1.26, df=29, p=0.22; Teosinte: t= - 0.76, df=30, p=0.25) or in N_2_O flux (B73: t=1.74, df=30, p=0.09; Teosinte: t= -0.07 df=30, p=0.95). In addition to this, we observed a dynamic pattern in N_2_O flux (mg m^-2^ min^-1^) across the season – where early season fluxes were similar but diverged by the end of the growing season (Fig. **4b**).

**Figure 4.**
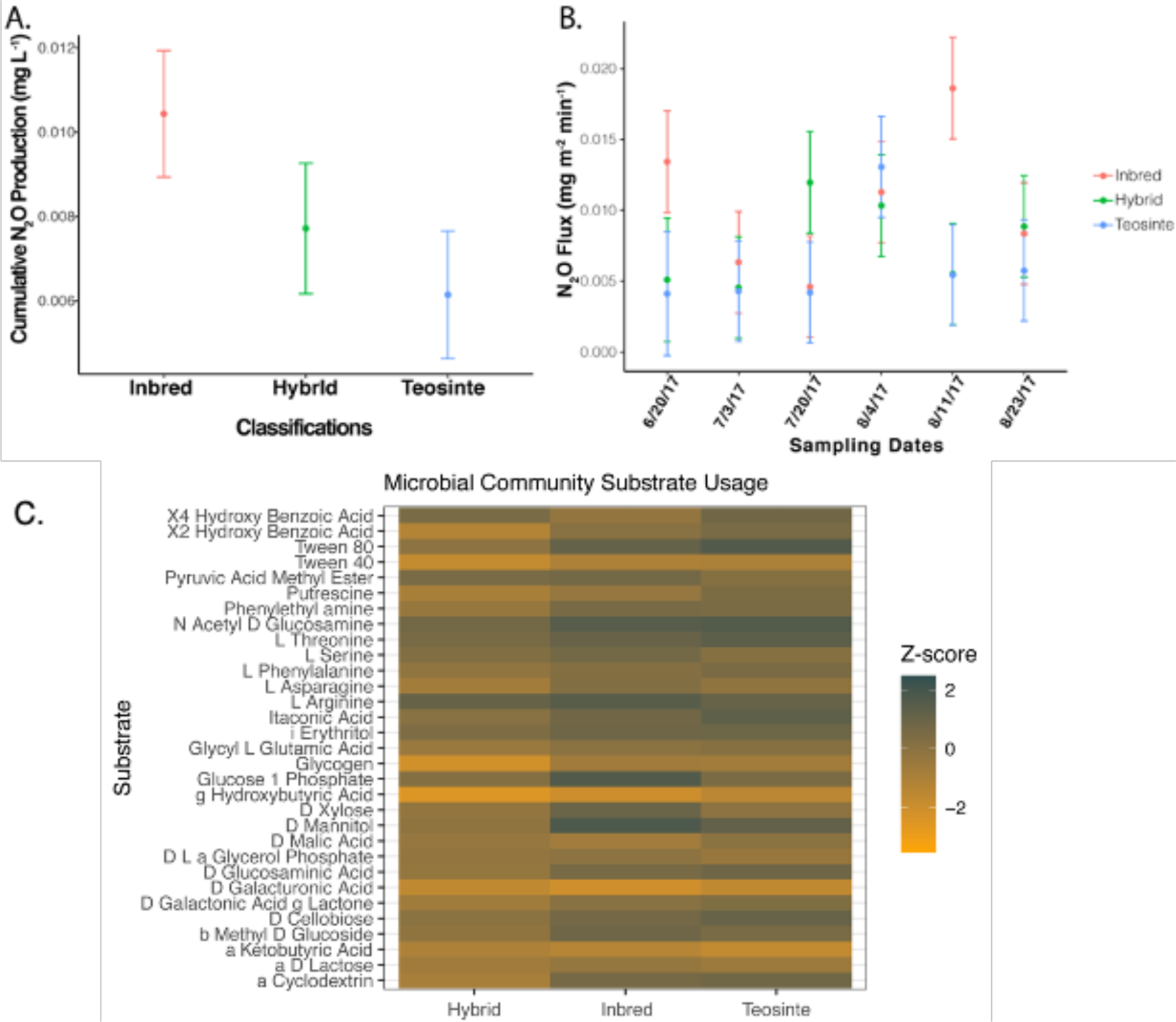
Cumulative N2O static flux chamber results over the growing season. **A.** LS means + 95% CI of cumulative N2O produced (mg L^-1^) averaged over the season comparing teosinte and B73 maize. **B.** LS means + 95% CI of N2O production flux (mg m^-2^ min^-1^) comparing teosinte and B73 maize displaying all timepoints. **C.** Carbon substrate usage (AWD) Z transformed compared among inbred B73, hybrid check1, and teosinte PI566677 (4 replicates per genotype) soil microbiomes measured using a BIOLOG-Eco microarray plates.

### Soil nutrient analysis

We found no significant difference in organic matter, estimated nitrogen release, cation exchange capacity, pH, sodium, iron, and boron between maize, hybrid, and teosinte plots. Teosinte plots had significantly higher levels of nitrate, phosphorus, potassium, sulfur, and manganese compared to inbred and hybrid maize plots (Wald’s test: p < 0.05; Table **S11).** Inbred maize plots had higher levels of buffer pH, copper, and calcium compared to teosinte and hybrid plots (Wald’s test: p < 0.05; Table **S11.)**

### Carbon substrate utilization

While not significant, we found that plant classification could potentially explain a substantial amount variation in microbial carbon substrate utilization (PERMANOVA: DF=2, R^2^=0.25, *p=*0.10). Between inbred B73, hybrid check1, and teosinte PI566677 genotypes, we found that teosinte root zone microbiomes had greater overall levels of substrate utilization compared to inbred and hybrid microbial communities (Fig. **4c**). We found no significant differences in utilization of individual substrates between inbred maize and teosinte microbial communities, 3 significant differences in substrate utilization between inbred and hybrid maize, and 2 significant differences in substrate utilization between hybrid and teosinte microbiomes (ANOSIM, *p*<0.05). Inbred-hybrid differences include glycogen, D-cellobiose, and L-serine. Teosinte-hybrid differences include L-phenylalanine and tween-80.

### Root Zone microbiome variation across the field

In this field experiment, we identified 37,596 different 16S rRNA operational taxonomic units (OTUs, 97% similarity, (rarefied to 100,000 reads per sample), and 2236 fungal OTUs (rarefied to 10,000) were identified from the ITS2 region (Table **S2**).

Within the prokaryotic community (based on 16S rRNA gene sequences), we found that plant genotype, genotype × time interactions, the location of the block, and sampling time explained 74% of the variation within the microbiome; 26% of the variation within the microbial community was unexplained (PERMANOVA, DF=26, p<0.001; Fig. **1**, S**7**; Table **S10**). In total, 34% of the variation in the prokaryotic microbiome was explained by plant genetics; 18% of this 34% variation was independent of temporal effects while 16% was highly linked to the time of sampling (genetics × sampling time). Interestingly, 20% of the variation in the soil microbial community was explained by block location alone. This would mean that across time, 20% of the microbiome was unchanged across the season. Sampling time (independent of genotype) explained 12% of the variation within the microbiome. Roughly, these results suggest that plant genetics explained about a third of the variation in the agroecosystem microbiome. Respectively, spatial and temporal effects seem to explain a third of the variation within the microbiome. Finally, an additional third of variation within the soil microbiome was unexplained.

Across plant classification (inbred, hybrid, teosinte), the most divergent genotype points showed the greatest differences in microbiome recruitment (Fig. S**7-8**; Table **S10.1-3**). Specifically, teosinte and hybrid maize treatments have the strongest effect on the composition of the soil microbiome agroecosystem. Teosinte root zone soils contained greater relative abundance of *Actinobacteria* and *Proteobacteria* (specifically, *Actinomycetales, Burkholderiales)* and less *Acidobacteria* (*iii1-15*, *Solibacteres)* compared to modern maize (Fig. **S9-10**). Additional analysis was carried out within plant category (i.e., within inbred, within hybrid, within teosinte), and inbred maize was the only category where genotype did not significantly contribute to differential microbiome recruitment.

Fungal communities showed similar results as the prokaryotic communities, except for notably weaker effects of space. This may indicate that fungi are less dispersal-limited than bacterial communities (Table **S10**).

### Relationship between the microbial community and N-cycling function

To further understand the differential contribution of the root zone microbiome to the potential function of a soil sample, we used weighted gene correlation network analysis (WGCNA) (Langfelder and Horvath, 2008) to identify four unique co-correlated clusters of OTUs (modules) with a significant response to potential function (3 modules that were correlated to potential nitrification, and one module that was correlated to overall denitrification, Fig. **5**). WGCNA taxa Module 2 was positively correlated to nitrification (r=0.20, p<0.001), while two modules were negatively correlated to nitrification (Module 4: r=-0.25, p<0.001; Module 7: r=-0.23, p<0.001). Module 2 contained 129 OTUs and was dominated by the presence of *Acidobacteria.* Interestingly, the second most dominant phylum in this module – *Cholorflexi* – was recently shown to have the ability to carry out nitrification (Spieck et al., 2020). Module 4 contained 290 OTUs and Module 7 contained 38 OTUs, both modules were dominated by *Actinobacteria.* Module 6 was correlated to the rate of overall denitrification (r=0.17, p<0.001), contained 26 OTUs, and was dominated by *Actinobacteria*.

**Figure 5.**
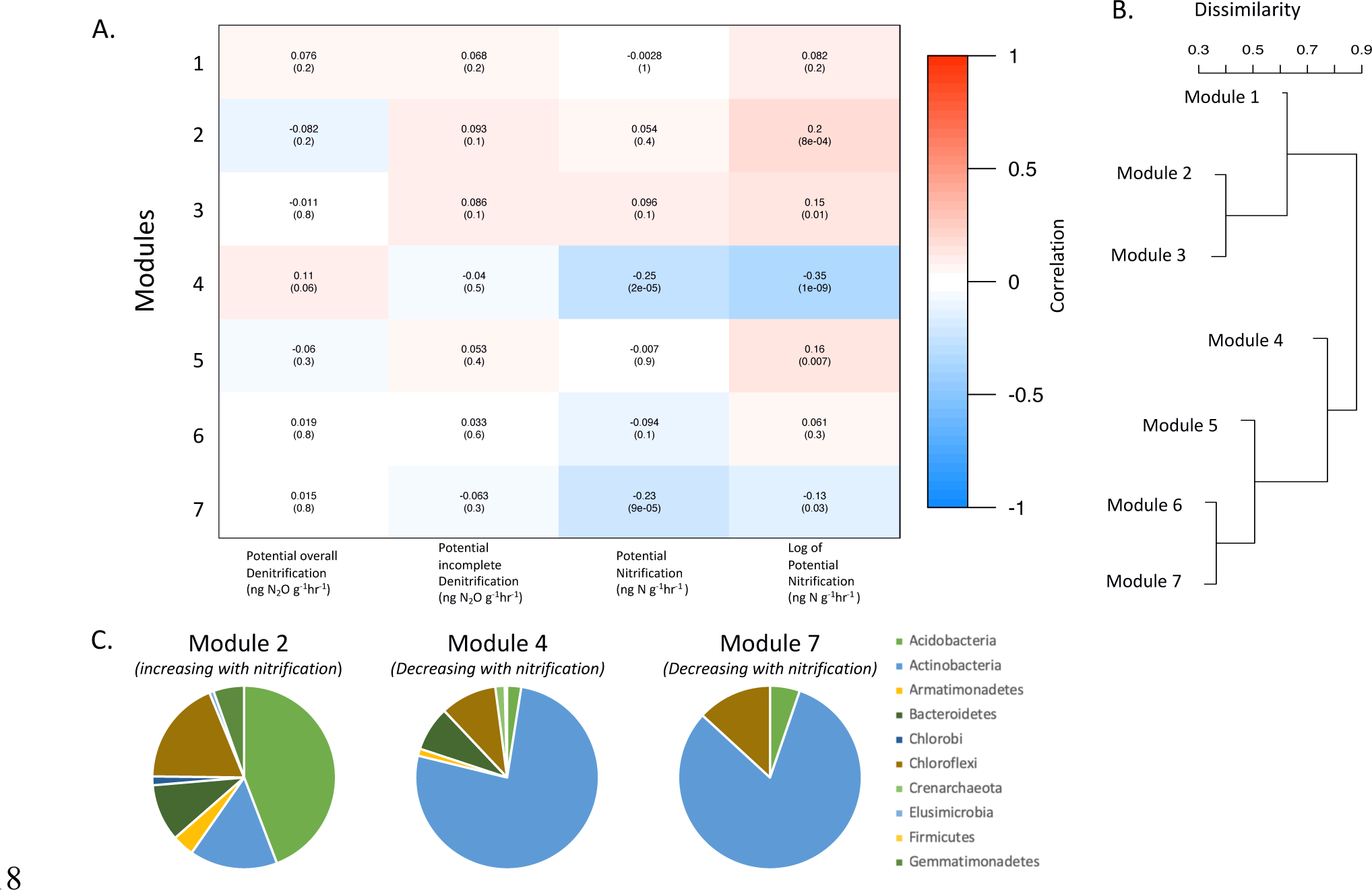
WGCNA results between potential denitrification and potential nitrification and microbial community composition. WGCNA starts by clustering microbial OTUs into modules of highly correlated taxa (based on abundance). These modules are then regressed against our explanatory factor (here that is denitrification and nitrification. **A.** Strength and significance of correlation between microbial modules and nitrogen cycling function. **B.** Correlations among microbial modules generated in the clustering process. **C.** Phylum-level composition of modules that were significantly correlated to changes in potential nitrification and denitrification.

## Discussion

### Microbial N-cycling processes modulated by Genetics

Across a variety of environmental conditions and plant species, it has been shown that genetic variation within a plant population can consistently shape the rhizosphere microbiome (Peiffer et al., 2013; Xu et al., 2018; Deng et al., 2021). Additionally, previous studies have also shown that changes in rhizosphere microbiome functional groups resulting from host plant genetics are important to ecosystem processes (i.e., carbon and nitrogen cycling) (Bulgarelli et al., 2015; Mwafulirwa et al., 2021a, 2021b). Furthermore, for decades researchers have known that plant carbon contributions to soils play a significant role in modulating microbial nitrogen transformations (Haller and Stolp, 1985; Qian et al., 1997). However, there is a lack of data showing how host genotype-specific changes in microbiome functional groups influence ecosystem processes. This field study demonstrates that the effect of plant genotype extends to modulating functions of the root zone microbiome. Specifically, we observed that the plant genotype influenced the recruitment of functional groups related to nitrification and denitrification along with the potential rates of those ecosystem processes (Fig **1-3**). These findings highlight the feasibility of breeding for plant phenotypes to influence N-cycling microbes and functions as a microbiome-associated phenotypes (MAPs) (Oyserman et al., 2018). Ultimately, our results suggest that we can select genetic haplotypes linked to MAPs within populations of agricultural cultivars to promote sustainable ecosystem processes within agroecosystems.

Our study demonstrates that genetic variation within *Zea mays* plays a significant role in both the assembly of the microbiome and the nitrogen cycling capability of the community, even in a stochastic field setting. Our most genetically heterogenous treatments (wild and domesticated hybrid) had the strongest effects on the composition and function of the microbiome. We hypothesized that the weak genotypic effects of inbred maize were driven by inbreeding in a low-stress, high nutrient environment, resulting in the alteration or loss of complex ecologically important traits, such as alteration of root architecture, microbiome recruitment, and plant exudate composition (Smýkal et al., 2018; Favela et al., 2021, 2022; Ren et al., 2022). Our hybrids showed distinct patterns of microbial community recruitment from their inbred parents, suggesting that heterosis plays a role in recruitment and function of the root zone microbiome. Interestingly, others have found that the expression of heterosis for root biomass and germination can be modulated by the presence of a soil microbiome (Wagner et al., 2020, 2021). Understanding the genetic basis of MAP heterosis is critical for designing efficient breeding programs for optimizing soil microbiome functions. Furthermore, we found that teosinte genotypes had the strongest influence on the activity of soil microbial taxa. We hypothesize that these differences in the teosinte microbiome were largely driven by a diverse set of belowground phenotypes (Gaudin et al., 2011), which likely includes alterations in exudate production, composition of metabolites, and root morphology. These results suggest that selective reincorporation of traits important to N-cycling would be key MAPs for “rewilding” modern breeding programs for sustainability (Perino et al., 2019; Razzaq et al., 2021). Importantly, this field study confirms that N-cycling functions, as well as composition of the rhizosphere microbiome, are responsive to genetic variation within the plant host.

### Diversity in inhibition Nitrification and Denitrification across Maize genotypes

We observed that teosinte and hybrid maize had the capacity to support lower activities of nitrifiers, while inbred maize stimulated their activity (Fig. **2a, 3a**). These results suggest that teosinte and hybrids may have a previously unrecognized mechanism to biologically suppress nitrification activity, an ability that has been weakened in maize. Biological nitrification inhibition (BNI) has been seen across a variety of different grass species (particularly in wild varieties) (Coskun et al., 2017a, 2017b) with recent work showing that modern maize does have some capacity for BNI (Otaka et al., 2022). BNI is a MAP by which plants exude secondary plant metabolites into the rhizosphere to inhibit nitrifying microbes which would typically compete with the plant for ammonium (Coskun et al., 2017b; Subbarao and Searchinger, 2021). These BNI mechanisms may have deteriorated through domestication bottlenecks, plant breeding, and introduction of modern agronomic practices, as it confers no benefit and may even be costly in the nitrogen-replete conditions of modern agroecosystems (as seen in wheat), and a parallel to what has been reported for nutritional symbioses with nitrogen-fixing microorganisms in legumes (Smercina et al., 2019; Favela et al., 2021; Subbarao et al., 2021). Investigating whether wild teosinte characteristics that confer improved BNI can be reincorporated into modern maize maybe of interest to future breeders, as nitrification can result in major losses of N fertilizers through leaching and GHG production (Nair et al., 2020; Subbarao et al., 2021). Furthermore, teosinte germplasm may be useful for bioprospecting novel BNI compounds that can be applied directly to agricultural fields (Kanchiswamy et al., 2015). Evaluating the vast diversity of maize cultivars for the capacity to carryout BNI using genetic information gleaned from teosinte is a valuable first step toward incorporating and improving these novel MAPs in agriculturally viable germplasm.

Interactions of plant genotypes with denitrifying microorganisms followed a similar pattern to nitrifiers, except with considerably more variation. Over the growing season, genotype and cultivar classification played a significant role in shaping potential denitrification activity (Figs. **2b-c, 3b-c**). These results were surprising, as maize is typically grown in aerobic soils and denitrification is an anaerobic process. Denitrifiers are often facultative anaerobes and may not be active under field conditions, only detected under the ideal conditions in the potential assay. Interestingly, teosinte appears to support lower potential incomplete denitrification (N_2_O) and overall denitrification (N_2_O + N_2_) leading to the hypothesis that teosinte contains biological denitrification inhibition traits (BDI) not previously reported. Biological denitrification inhibition is hypothesized to evolve in plants as a mechanism to compete with denitrifying microbes for soil nitrates. As with BNI, the mechanisms underlying BDI typically involve the release of phytochemicals that directly interfere with the enzymes carrying out N-metabolism (Bardon et al., 2016). Previous work in rice, a crop grown in anaerobic soil, has also shown that genetic variation exists within the rice species to allow inhibition of denitrifier activities (Ishii et al., 2011; Ding et al., 2019). Compared to BNI research, BDI work is still in early stages with only a single class of metabolites, procyanidins, being shown to mediate BDI (Galland et al., 2019). Procyanidins inhibits denitrification by acting as a allosteric inhibitor of nitrate reductase (Bardon et al., 2016). Interestingly, a considerable amount of work has focused on a maize depolarized procyanidin (cyanidin) and anthocyanin (a glucoside cyanidin), showing that maize genotypes have considerable variation in cyanidin and anthocyanin production (Sharma et al., 2011; Paulsmeyer et al., 2017). While not quantified in this study, the BDI differences observed here among maize genotypes could potentially be related to differences in cyanidin and anthocyanin exudation in the rhizosphere. It has been shown that maize cultivars are able to produce anthocyanin in the root and that closely related species like *Sorghum* produce anthocyanins at the root tip (Tselas et al., 1979; Hawes et al., 1998). It is possible that these rhizosphere cyanidins and anthocyanin are acting as anti-reductants, which is known to occur at low pH (Becker, 2016), and are competitive inhibitors of denitrification or allosteric inhibitors like procyanidins. Further research needs to be done to determine if maize anthocyanins have the capacity to inhibit denitrification. These BDI MAPs are valuable, as they present a mechanism to limit denitrification (and thereby GHG production) if reintroduced to the agroecosystem.

### Outcomes of N-cycling alteration by Plants

From a sustainability perspective, this study highlights a potential avenue to reduce agricultural N losses and GHG emissions generated by soil microorganisms. N_2_O static chamber results (Fig. **4a-b**) provide support that these potential nitrification and denitrification assays may, in fact, reflect some aspect of ecosystem processes. It should be noted that these potential assays and N_2_O chambers, have their limitations (Nannipieri et al., 2018; Grace et al., 2020). Potential assays indicate that the maximum function of these N-cycling communities has changed in response to plant host in the root zone. While the N_2_O chambers shows that this may be related to some change in flux of N out of the system. This is an exciting finding, as agriculture is a major producer of N_2_O emissions (Vitousek et al., 1997; Reay et al., 2012), and these results may give us a foothold to potentially curb production of this potent GHG. While these N_2_O results presented are interesting, a major limitation of this study is that we only examined these cultivars in a single field and static flux chambers are known to be extremely variable (Waldo et al., 2019). Further support for our potential nitrification and denitrification assays translating to actual differences in field processes can be seen in our end of season soil nutrient analysis. From this analysis, we observed that teosinte plots maintained higher levels of nitrates (Table S11) compared to hybrid and inbred plots. These higher nitrate and nutrient levels were likely driven by the capacity to carry out BNI and BDI and ability to quell microbial pathways for nutrient loss. This enhanced ability to mine nutrients in teosinte may be facilitated by changes in root zone pH driven by carbon exudation, and root density. This conclusion is supported by our finding showing that teosinte plots have a lower buffer pH (residual or reactive pH) compared to hybrid and inbred maize. Alteration of soil buffer pH is likely underpinning many of the functional changes we observed in this study. In addition, we observed modest differences in microbial community carbon metabolism between plant classifications (Fig. **4c**) suggesting potentially different soil carbon cycling driven by the plant host. From these carbon enzyme assays, we found that teosinte-derived microbial communities have a heightened capacity to degrade phenylalanine, Tween-80, and oleic acid. Interestingly, phenylalanine is a critical precursor to the synthesis of flavonoids, while oleic acid is a precursor to linoleic acid, a previously characterized BNI molecule (Kaur-Bhambra et al., 2022). Perhaps this improved metabolism in teosinte-derived microbiomes was primed by previous exposure in the plant rhizosphere as exudates. Conversely, microbial communities associated with hybrid maize were better equipped to degrade carbohydrates like glycogen and cellobiose compared to the root zone microbiome of inbred maize. This result is interesting, as microbes differ in their ability to consume different photosynthates exuded into the rhizosphere (Fan et al., 2022), lending support to the hypothesis that differences we are observing across microbiomes may be caused by altered amount and composition of plant exudates. This hypothesis is further supported by work in barley showing that genotypes can vary in rhizodeposition-derived carbon and this variation shapes soil microbial mineralization (Mwafulirwa et al., 2016). The next steps toward incorporating potential nitrogen conservation MAPs into agricultural practices would be research to explore whether suppression of nitrogen transformations is consistent across a wide range of biogeographic environments.

The effects of seasonal phenology and *Zea* genotype × sampling time interaction over the growing season played a major role in microbiome recruitment and function. Maize has different nutrient requirements across the growing season, and these nutrients are extracted from the soil environment (Bender et al., 2013), so, along with genotype-specific biochemistry, plant growth and development influence interaction with the soil microbial community. In addition, previous studies have shown that the complexity of the microbiome is built through time (Shi et al., 2016; Emmett et al., 2020; Ajilogba et al., 2022). These temporal effects are important to consider, as they can dramatically influence the conclusions drawn about the interaction between plants and their microbiome. We observed that potential nitrification and potential denitrification were dependent on plant growth stage (Fig. **2**). Potential nitrification and BNI phenotypes, for example, seemed to peak in the middle of the season, coinciding with plant primary productivity (Fig. **2**). During peak growth, it appeared as if *Zea* was priming soils for release of plant-available nutrients. This process was shown to enhance the release of N (Phillips et al., 2011; Emmett et al., 2020), perhaps explaining the increase in nitrification. Furthermore, these temporal impacts highlight a limitation in this study, whereby we may be overestimating our genotype differences because of phenological growth between the cultivars. In addition, temporal growth patterns are likely interacting with the measurability of effects in the root zone as root density of the plant is anticipated to change through plant development(Chaparro et al., 2013; Vetterlein et al., 2020; Tkacz and Poole, 2021).. While not captured in this study, future work needs to delineate and find methods to capture growth and temporal variation belowground to meaningfully incorporate the microbiome and ecosystem processes into the agricultural setting (York et al., 2022).

### Relationship between Microbial diversity and N-cycling

We found that different microbial taxa were correlated with functional changes in the microbiome. WGCNA identified four modules of microbial taxa that were significantly associated with both changes of potential nitrification and denitrification (Fig. **5**). These results suggest that specific taxa and their interactions play a role in driving the function of the microbiome and that, to some degree, plants can influence the activities of specific microbial groups. Interestingly, we observed that modules positively correlated with higher potential nitrification rates were dominated by gram-negative bacteria within the phylum *Acidobacteria*. In contrast, those that were negatively correlated with nitrification were dominated by gram-positive bacteria within the *Actinobacteria* phylum. Overall denitrification was positively correlated to gram-positive bacteria (*Actinobacteria*). Perhaps, cell envelope differences play a role in how plants select taxa to join the microbiome or pH changes caused by nitrification and/or plant exudation facilitated the growth of *Acidobacteria*. Surprisingly, these modules of correlated OTUs were not dominated by known nitrifying taxa, suggesting that nitrification processes may be, in part, dependent on the metabolism of other microbial community members, that nitrification is controlled by the level of transcriptional regulation rather than nitrifying population size, nitrifiers or nitrifier ideal conditions are facilitating habitat alteration that is influencing microbiome structure, or that some other microbial interaction (i.e. predation, competition) is controlling nitrification (Baskaran et al., 2020; Spieck et al., 2020; Burian et al., 2021). This could be, in part, because the ammonia monooxygenase enzyme is readily inhibited by a variety of phytochemicals (Bedard and Knowles, 1989). For example, gram-negative bacteria that are positively correlated with nitrification may be breaking down phytochemicals that would otherwise inhibit ammonia monooxygenase, allowing nitrifiers to continue nitrification, while the gram-positive bacteria that are negatively correlated with nitrification may not be breaking down plant phytochemicals or may be producing some inhibitory compound. Determining how ecological interactions within the microbiome influence microbial functions is critical for understanding and predicting microbially-mediated ecosystem functions (Kuypers et al., 2018).

## Conclusion

The ability of plants to influence microbial functions in the rhizosphere likely evolved as a mechanism for nutrient retention, enhancing plant competition for available nutrients from the soil matrix (Philippot et al., 2013b; Delaux and Schornack, 2021; Lata et al., 2022). It is becoming increasingly clear that plant genetic variation (within and among species) modulates the activities of the soil-associated microbiome and that these alterations can impact soil biogeochemical functions (Falkowski et al., 2008; Morris et al., 2020). Identification of the plant genomic regions that direct recruitment of N-cycling microorganisms or modulation of their activities will enable progress on re-engineering the agroecosystem to become a minor contributor to N pollution (Johnson, 2006; Subbarao and Searchinger, 2021). This study adds to this growing body of work by showing that maize, an agronomically important crop, has genetic variation that contributes to alterations in the microbiomes and N-cycling function – potentially enough variation to breed and incorporate these extended phenotype ecosystem traits into modern hybrid cultivars. Integrating sustainability-related microbiome associated phenotypes into our agricultural systems is a way forward to address many agronomic challenges that society faces.

## Acknowledgements

We thank Savannah Henderson, and Sierra Raglin for assistance in the field and lab; Christopher Mujjabi and the Crop Sciences Research and Education Center with assistance field planting and maintenance; Mark Band at the University of Illinois Functional Genomics Center for assistance with the development of multiplex PCR using the Fluidigm system; and Chris Wright and the staff of the DNA Services Lab at the Roy J. Carver Biotechnology Center for assistance with DNA sequencing.

