## Supplemental Files for "Genetic variation exists within *Zea mays* to influence unsustainable nitrogen cycling microbiome function"

### Genetic variation within *Zea mays* alter microbiome assembly and nitrogen cycling function in the Agroecosystem

#### Supplemental Information and Materials

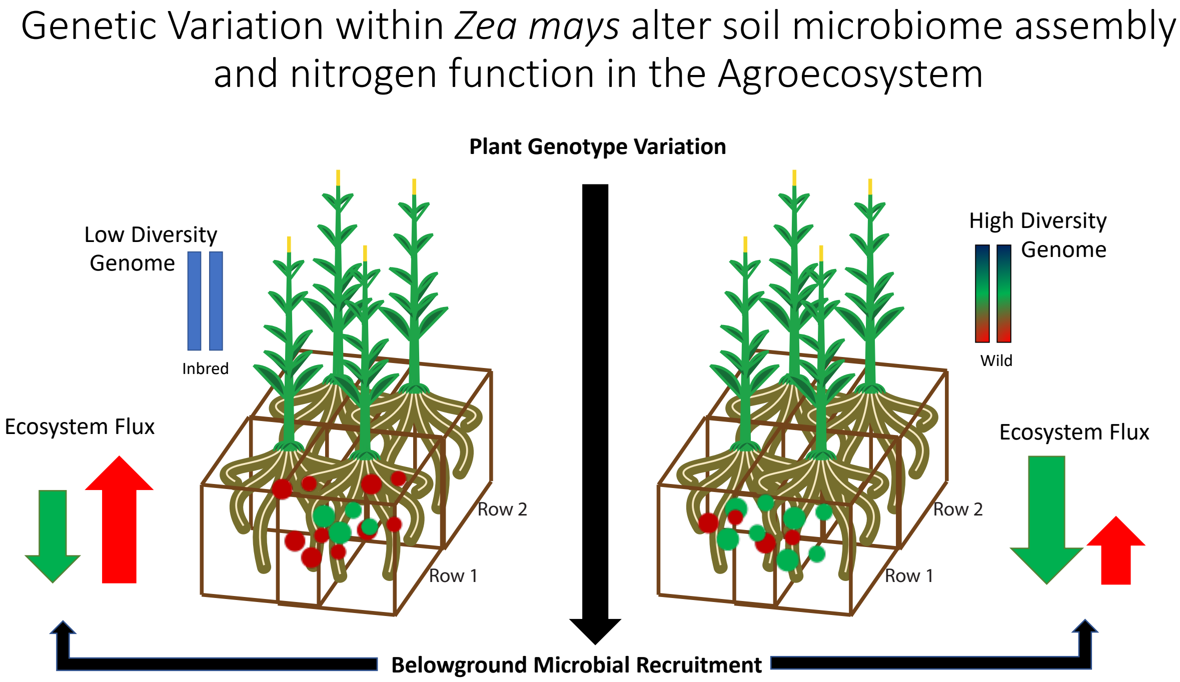

**Figure S1.** Graphical abstract displaying the underlying concept of the study. It has been shown that plant genotype plays a role in microbial requitement- here we were interested in understanding if genotype driven differential requitement of the rhizosphere microbiome will result in alterations to microbial mediated ecosystem functions. To assess this question, we used extremely contrasting genetic models: modern agricultural inbred of maize and wild teosinte progenitors. This gives us a potential ceiling to how much of a role plant genetic can play.

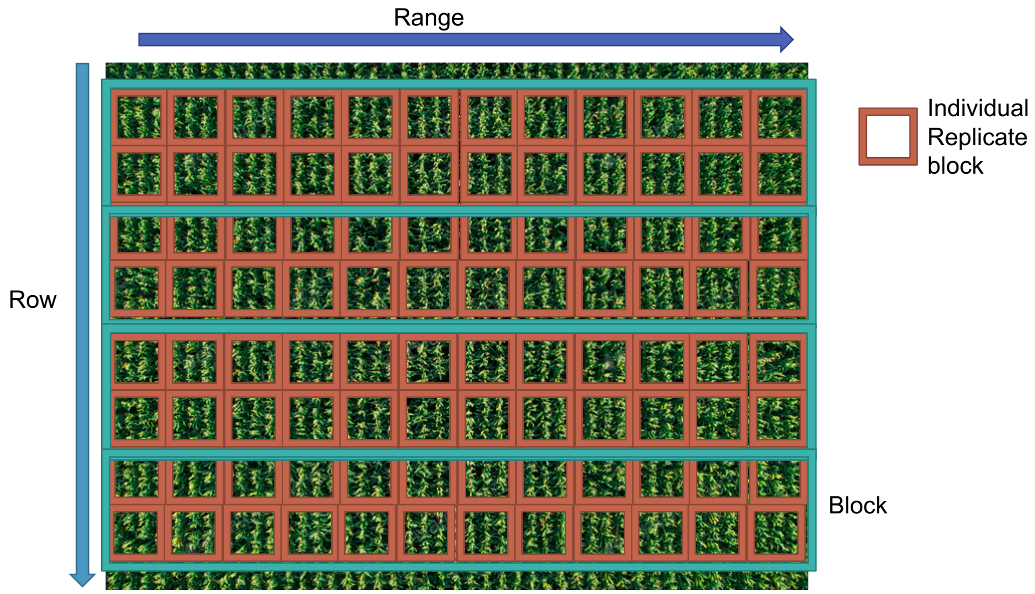

**Figure S2.** Physical structure of the field setting. Replicate rectangular blocks are colored in teal and include all the genotypes included in the study. Individual genotype blocks are colored in brown and are randomized within the block. Range and row factors were included in model as continuous factors to represent space within the field. This scheme allowed us to control for stochastic spatial effects within our experiment.

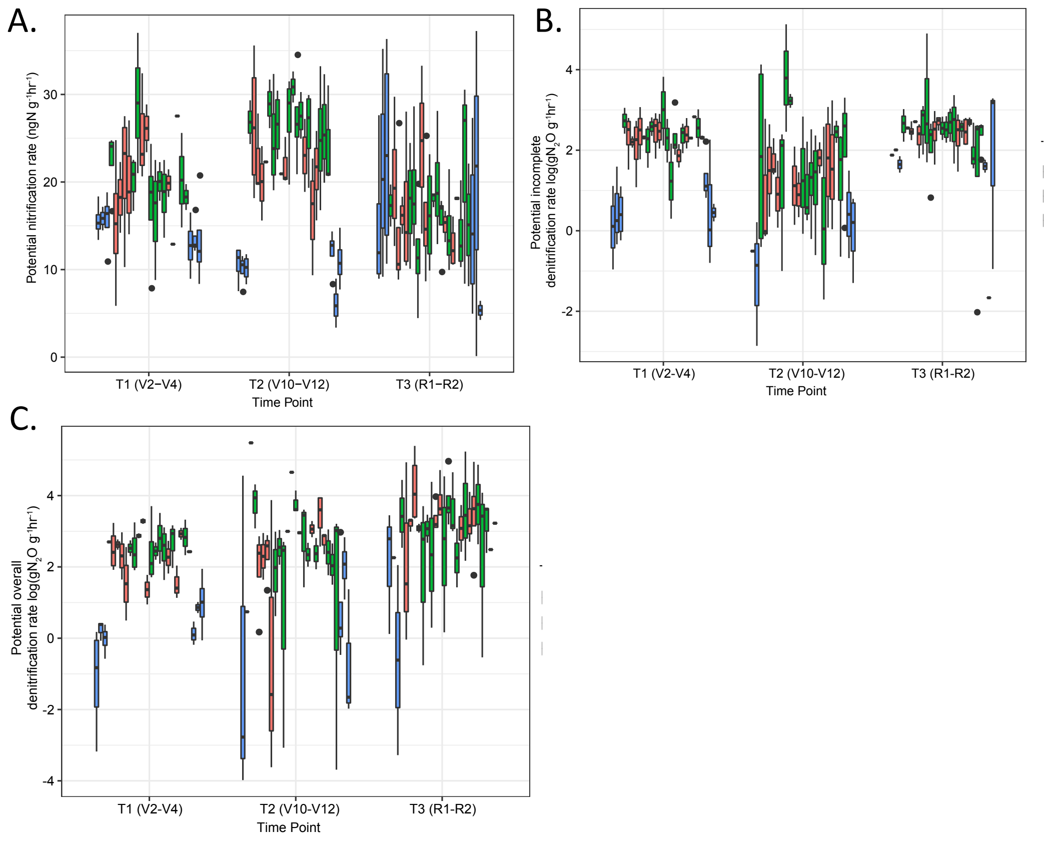

**Figure S3.** Rates of potential nitrification (ng N $g^{-1}$ $hr^{-1}$), potential incomplete denitrification (log (ng $N_2O$ $g^{-1}$ $hr^{-1}$)), and potential overall denitrification (log (ng $N_2O$ $g^{-1}$ $hr^{-1}$)) as influenced by plant genotype and classification. **A.** Potential nitrification rate (ng N $g^{-1}$ $hr^{-1}$) of all genotypes included in the study across our three sampling time points colored by plant classification. Teosinte genotypes were observed as having the lowest nitrification rates in the T1 and T2. No differences were present at the T3 sampling time point. **B.** Potential incomplete denitrification (log (ng $N_2O$ $g^{-1}$ $hr^{-1}$)) of all genotypes included in the study across our three sampling time points colored by plant classification. **C.** Potential overall denitrification (log (ng $N_2O$ $g^{-1}$ $hr^{-1}$)) of all genotypes included in the study across our three sampling time points colored by plant classification.

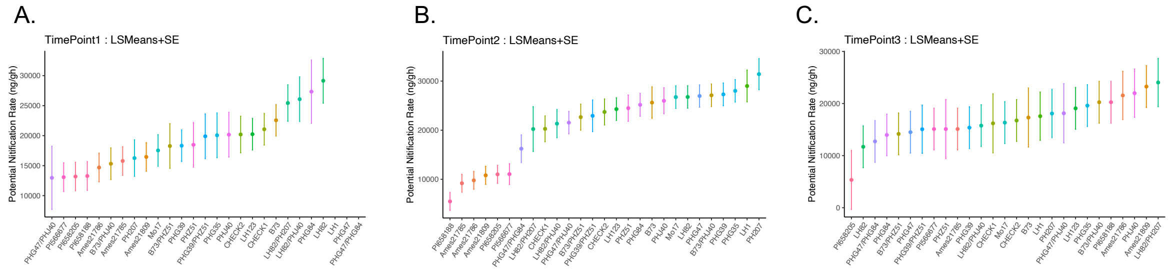

**Figure S4.** Potential nitrification of individual genotypes in study across different sampling timepoints. LS Means and Standard Error were calculated using ASREML-r. Supplemental statistics are presented in Table S7. **A.** Genotypic mean of potential nitrification at time point 1. **B.** Genotypic mean of potential nitrification at time point 2. **C.** Genotypic mean of potential nitrification at time point 3.

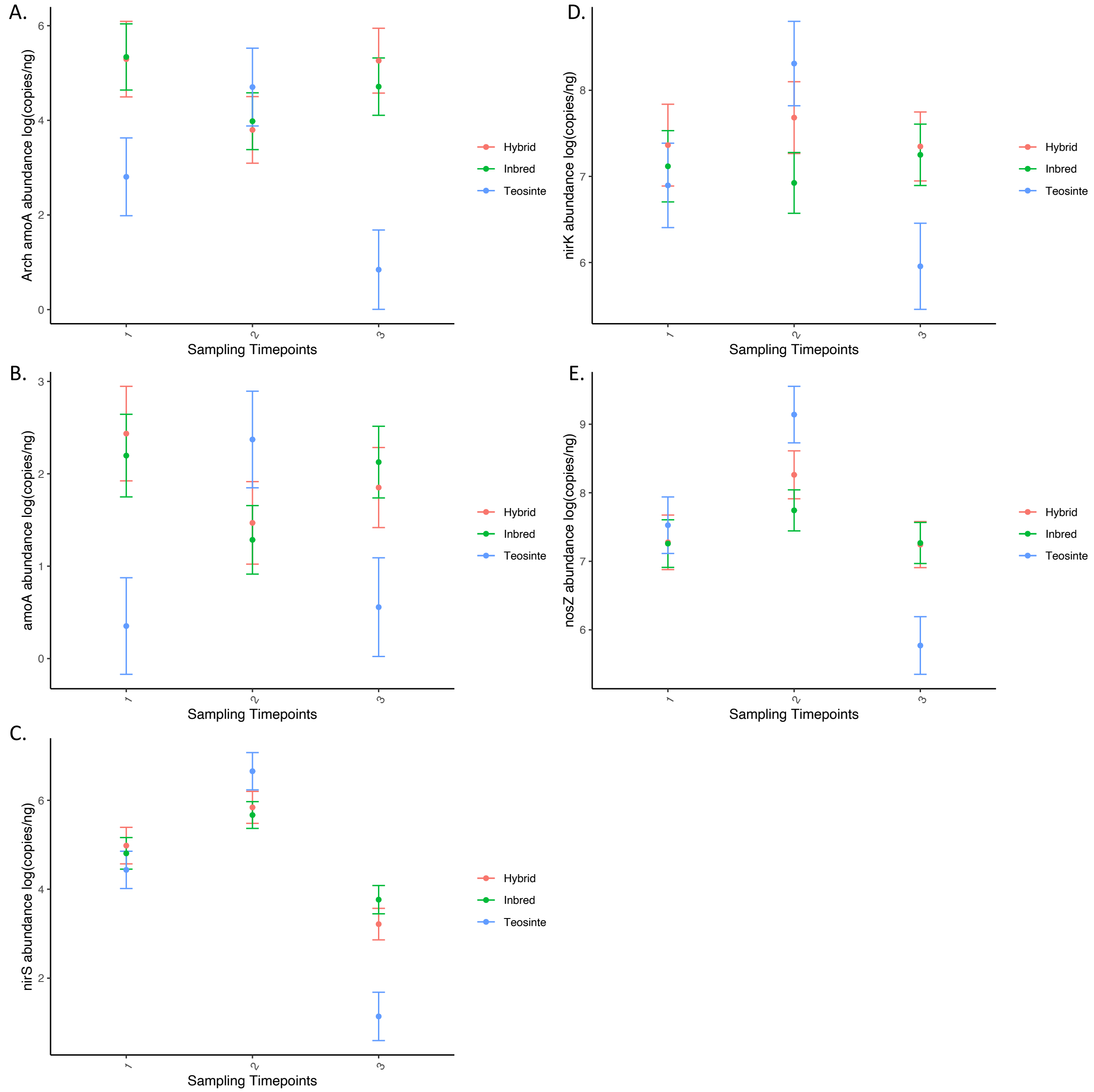

**Figure S5.** Quantitative PCR results for functional genes surveyed in the study across different timepoints over the season across plant classification. LS Means and Standard Error were calculated using ASREML-r. Supplemental Statistic are present in Table S6. **A.** Log of archaeal *amoA* qPCR abundance across timepoints. **B.** Log of bacterial *amoA* qPCR abundance across timepoints. **C.** Displays the log of *nirS* qPCR abundance across timepoints. **D.** Log of *nirK* qPCR abundance across timepoints. **E.** Log of *nosZ* qPCR abundance across timepoints.

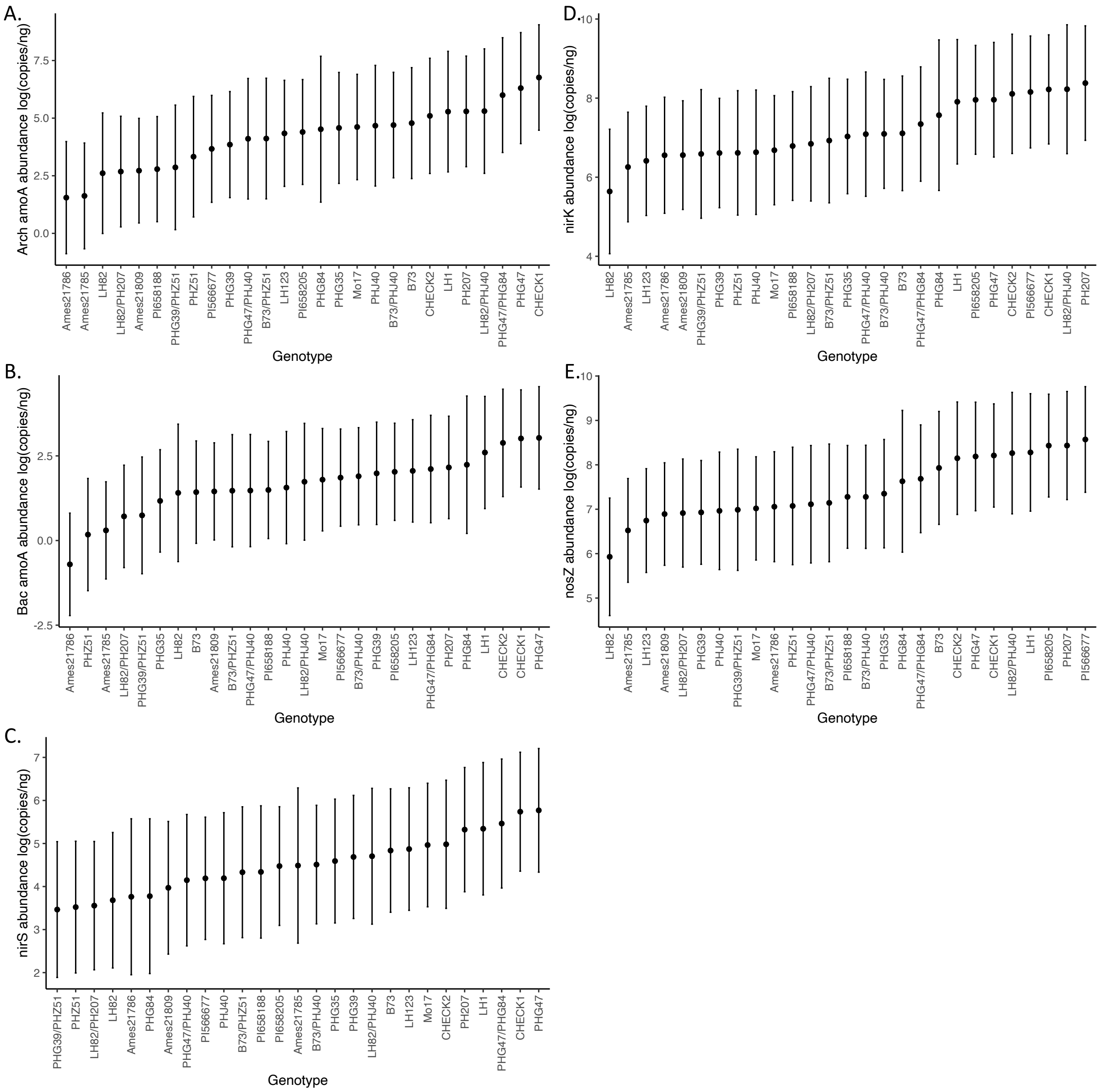

**Figure S6.** Quantitative PCR Results for functional genes surveyed in the study averaged cross different timepoints displayed as genotypic means. LS Means and Standard Error were calculated using ASREML-r. Supplemental Statistic are present in Table S6. **A.** Log of archaeal *amoA* qPCR abundance across genotypes. **B.** Log of bacterial *amoA* qPCR abundance across genotypes. **C.** Log of *nirS* qPCR abundance across genotypes. **D.** Log of *nirK* qPCR abundance across genotypes. **E.** Log of *nosZ* qPCR abundance across genotypes.

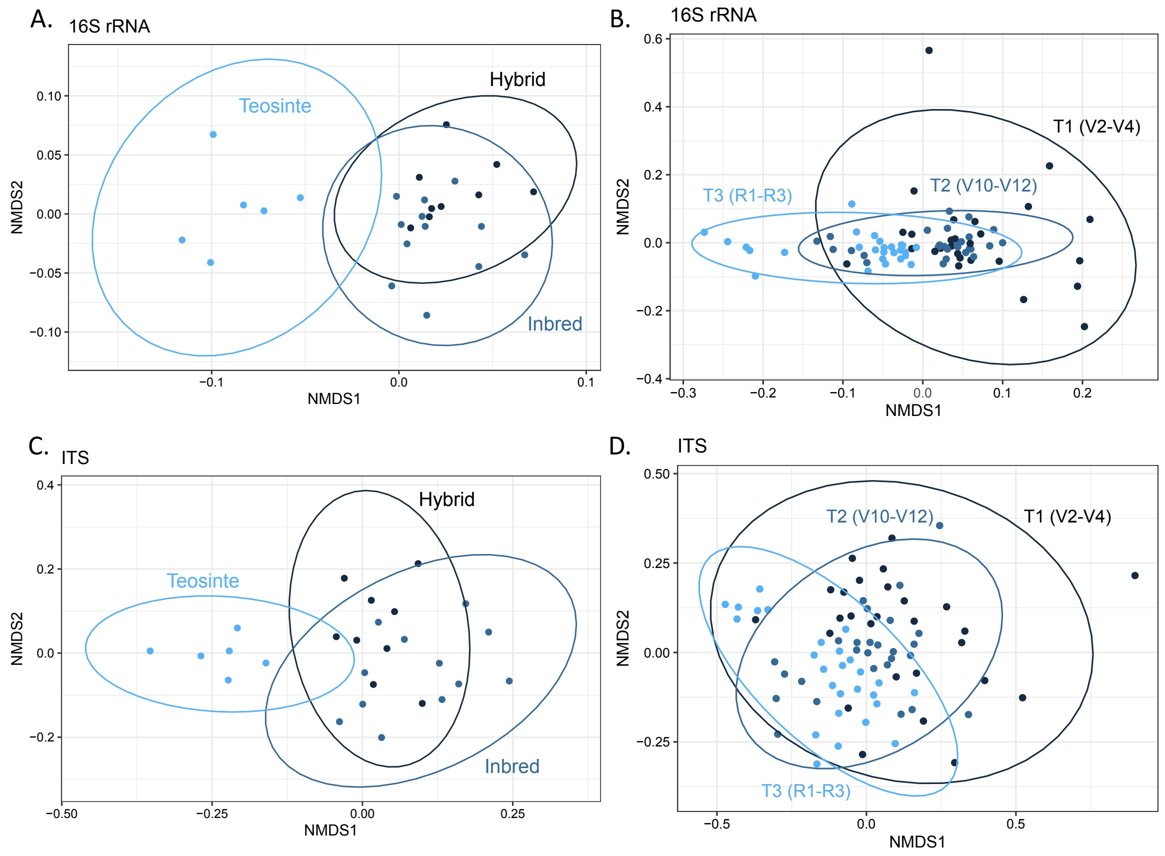

**Figure S7.** NMDS ordinations based on Bray-Curtis dissimilarity among prokaryotic 16S rRNA (**A-B**) and fungal ITS (**C-D**) displaying that genetic (**A, C**) and temporal (**B, D**) effects are driving changes in composition of the root zone microbiome. (**A, C**) shows that inbred, hybrid, and teosinte maize lines host different microbial taxa in the root zone under the same environmental conditions. Each point represents a genotypic mean (within mean n= 12) of the microbial community across the three different sampling time points. (**B, D**) highlights the temporal effects of the rhizosphere microbiome timepoint 1 (young plants V2-V4), 2 (Fast growing approaching flowering V10-V12), 3 (Reproductive R1-R3). (Within mean n=4). Statistics for ordinations present in Table S10.

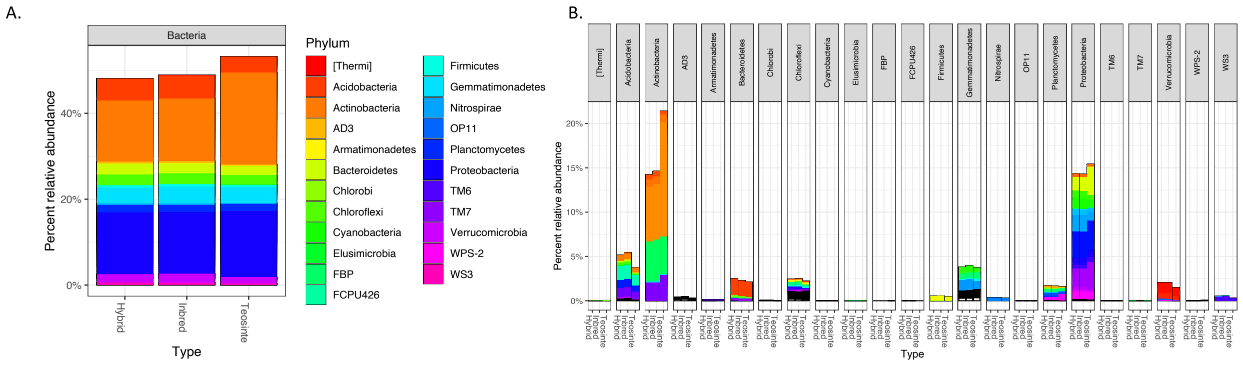

**Figure S8.** OTUs that differed significantly based on DESeq2 analysis of microbiomes associated with teosinte and inbred maize. **A.** Relative abundance of bacterial OTUs colored by phylum. The most contrasting differences were seen in *Acidobacteria*, *Actinobacteria,* and *Proteobacteria as seen in S9*. **B.** This panel shows the same relative abundance data as in A but faceted by phylum and colored by taxonomic classification to improve clarity. These taxa were major contributors to the microbiome differences displayed in Figs. S6-7

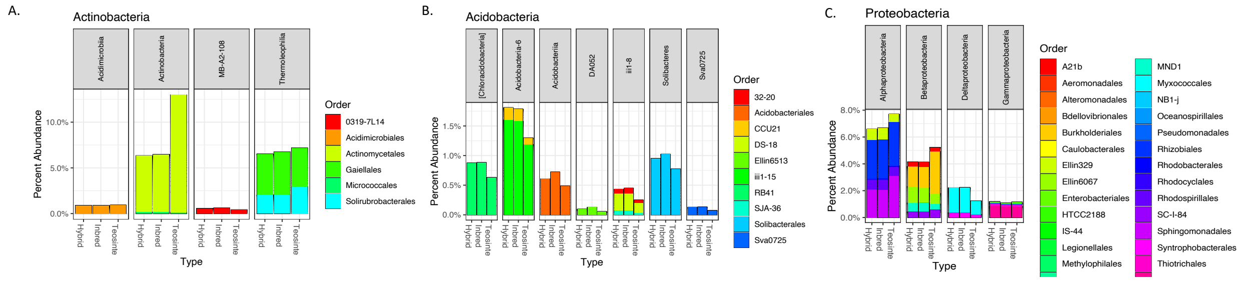

**Figure S9.** Differentially abundant OTUs based on DESeq2 determined by teosinte and inbred maize comparison. **A.** OTUs within order *Actinobacteria* that differ among teosinte and inbred maize. **B.** OTUs within order *Acidobacteria* that differ among maize classifications. **C.** OTUs within order *Proteobacteria* that differ among maize classifications. These OTUs contribute to differences in microbiome assemblages displayed in Figures S7.

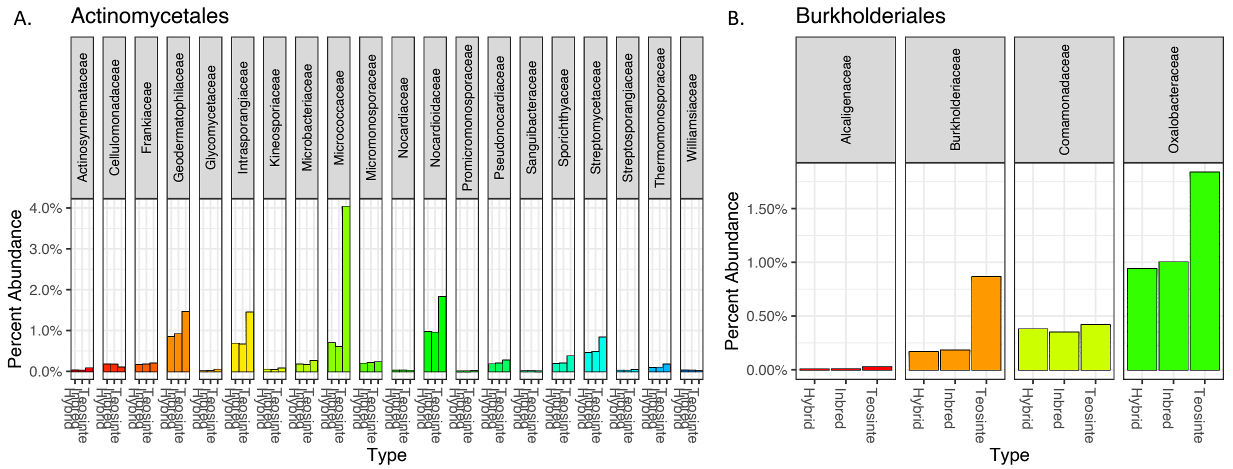

**Figure S10.** OTUs with significantly different relative abundance, based on DESeq2 analysis comparing the microbiomes of teosinte and inbred maize . These taxa were major contributors to the microbiome differences displayed in Figs. S6-7. **A.** OTUs within the *Actinomycetales* that were significantly different between teosinte and inbred maize. **B.** OTUs within the order *Burkholderiales* that were significantly different between teosinte and inbred maize.

#### Supplemental Information: Tables

**Tables S1.** Accession table of lines grown in this experiment. Hybrid genotypes were generated from previous crosses in the field.

| Genotypes | Classification | Accession # |
|---|---|---|
| B73 | Inbred | PI 550473 |
| LH1 | Inbred | PI 644101 |
| LH123 | Inbred | PI 601079 |
| LH82 | Inbred | PI 601170 |
| Mo17 | Inbred | PI 558532 |
| PH207 | Inbred | PI 601005 |
| PHG35 | Inbred | PI 601008 |
| PHG39 | Inbred | PI 600981 |
| PHG47 | Inbred | PI 601318 |
| PHG84 | Inbred | PI 601320 |
| PHJ40 | Inbred | PI 601321 |
| PHZ51 | Inbred | PI 601322 |
| B73/PHJ40 | Hybrid | NA |
| B73/PHZ51 | Hybrid | NA |
| CHECK1 | Hybrid | NA |
| CHECK2 | Hybrid | NA |
| LH82/PH207 | Hybrid | NA |
| LH82/PHJ40 | Hybrid | NA |
| PHG39/PHZ51 | Hybrid | NA |
| PHG47/PHG84 | Hybrid | NA |
| PHG47/PHJ40 | Hybrid | NA |
| PI566677 | *Teosinte: Z. M. Mexicana* | PI566677 |
| PI658188 | *Teosinte: Z. M. Mexicana* | PI658188 |
| PI658205 | *Teosinte: Z. M. Mexicana* | PI658205 |
| Ames21809 | *Teosinte: Z. M. Parviglumis* | Ames21809 |
| Ames21785 | *Teosinte: Z. M. Parviglumis* | Ames21785 |
| Ames21786 | *Teosinte: Z. M. Parviglumis* | Ames21786 |

**Table S2.** Primers used in amplicon sequencing and qPCR characterization of rhizosphere microbial community.

| Target | Encodes | Primer Name | Sequence | Reference |
|---|---|---|---|---|
| *16S rRNA* | Ribosomal RNA | 515F | 5'-GTGYCAGCMGCCGCGGTAA-3' | Fierer et al. 2011 |
| *16S rRNA* | Ribosomal RNA | 806R | 5'-GGACTACVSGGGTATCTAAT-3' | Fierer et al. 2011 |
| *ITS* | Internal Transcribed Spacer | ITS1F | 5'-TTCGTAGGTGAACCTGCGG-3' | White et al. 1990 |
| *ITS* | Internal Transcribed Spacer | ITS4R | 5'-TCCTCCGCTTATTGATATGC-3' | White et al. 1990 |
| bacterial *amoA* | Ammonia Monooxygenase | amoA-1F | 5'-GGGGTTTCTACTGGTGGT-3' | Oakley et al. 2005 |
| bacterial *amoA* | Ammonia Monooxygenase | amoA-2R | 5'-CCCCTCKGSAAAGCCTTCTTC-3' | Oakley et al. 2005 |
| archeal *amoA* | Ammonia Monooxygenase | CrenamoA23f | 5'-ATGGTCTGGCTWAGACG-3' | Francis et al. 2005 |
| archeal *amoA* | Ammonia Monooxygenase | CrenamoA616r | 5'-GCCATCCATCTGTATGTCCA-3' | Francis et al. 2005 |
| Typical *nosZ* | Nitrous oxide reductase | nosZ1F | 5'-WCSYTGTTCMTCGACAGCCAG-3' | Henry et al. 2006 |
| Typical *nosZ* | Nitrous oxide reductase | nosZ1R | 5'-ATGTCGATCARCTGVKCRTTYTC-3' | Henry et al. 2006 |
| *nirK* | Nitrite Reductase | nirK876 | 5'-ATYGGCGGVCAYGGCGA-3' | Henry et al. 2004 |
| *nirK* | Nitrite Reductase | nirK1040 | 5'-GCCTCGATCAGRTTRTGGTT-3' | Henry et al. 2004 |
| *nirS* | Nitrite Reductase | nirSCd3aF | 5'-AACGYSAAGGARACSGG-3' | Kandeler et al. 2006 |
| *nirS* | Nitrite Reductase | nirSR3cd | 5'-GASTTCGGRTGSGTCTTSAYGAA-3' | Kandeler et al. 2006 |
| *norB* | Nitric Oxide Reductase | cnorB2F | 5'-GACAAGNNNTACTGGTGGT-3' | Braker et al. 2003 |
| *norB* | Nitric Oxide Reductase | cnorB6R | 5'-GAANCCCCANACNCCNGC-3' | Braker et al. 2003 |

**Tables S3.** Quality of amplicon sequencing for all genes in this study.

| Amplicon | Raw Reads | Quality Filtered Reads | Rarefaction Level |
|---|---|---|---|
| **16S rRNA gene** | 79,484,164 | 74,085,411 | 100000 |
| **fungal ITS** | 8,531,066 | 7,766,289 | 10000 |
| **bacterial *amoA*** | 1,613,336 | 1,478,539 | 337 |
| **archaeal *amoA*** | 3,561,327 | 3,257,457 | 2500 |
| ***nirK*** | 23,066,482 | 23,143,193 | 1948 |
| ***nirS*** | 6,239,375 | 5,661,656 | 1164 |
| ***nosZ*** | 7,071,538 | 9,306,149 | 2202 |

**Table S4.** Functional gene summary of amplicon functional composition gene and qPCR abundance data from PerMANOVA and ASREML statistical analysis. S4.1 displays the genotype summary. S4.2 displays the plant classification summary. Full statistical model presented in S5-6.

**Table S4.1.** Overview of statistical analysis for genotypes effects on different functional genes.

| Ecosystem Process | Gene | Significant Composition Across Genotype ($p<0.05$) | Significant Composition Genotype:Time Interaction ($p<0.05$) | qPCR Abundance Significance in Relation to Genotype ($p<0.05$ ) | qPCR Abundance Significance Genotype:Time Interaction ($p<0.05$) |
|---|---|---|---|---|---|
| Nitrification | Bacterial *amoA* | Non-Significant | **Significant** | Non-Significant | Non-Significant |
| Nitrification | Archeal *amoA* | **Significant** | **Near-Significant (p=.08)** | Non-Significant | Non-Significant |
| Denitrification | *nirK* | **Significant** | Non-Significant | Non-Significant | **Near-Significant (p=.06)** |
| Denitrification | *nirS* | **Significant** | Non-Significant | Non-Significant | **Near-Significant (p=.06)** |
| Denitrification | *nosZ* | **Significant** | Non-Significant | **Near-Significant (p=.08)** | **Near-Significant (p=.07)** |

**Table S4.2** Overview of statistical analysis for classification (inbred, hybrid, teosinte) effects on different functional genes.

| Ecosystem Process | Gene | Significant Composition Across Type ($p<0.05$) | Significant Composition Class:Time Interaction ($p<0.05$) | qPCR Abundance Significance in Relation to Class ($p<0.05$ ) | qPCR Abundance Significance Class:Time Interaction ($p<0.05$) |
|---|---|---|---|---|---|
| Nitrification | Bacterial *amoA* | Non-Significant | **Significant** | Non-Significant | **Significant** |
| Nitrification | Archeal *amoA* | **Significant** | **Significant** | Non-Significant | Non-Significant |

| | | | | | |
|---|---|---|---|---|---|
| Denitrification | *nirK* | **Significant** | Non-Significant | Non-Significant | **Significant** |
| Denitrification | *nirS* | **Significant** | Non-Significant | Non-Significant | **Significant** |
| Denitrification | *nosZ* | **Significant** | **Significant** | **Significant** | **Significant** |

**Table S5.** Functional Gene Summary of amplicon functional from PerMANOVA models. 999 permutations were used in analysis. All factors in the model were run as fixed effect: ***Microbial Community Matrix = Genotype (or Classification) x Sampling Time + Row x Range + Residuals.*** Interactions between plant classification (inbred, hybrid, teosinte) and time were incorporated to understand plant growth changes, while row and range interactions were used to understand spatial effects. Row and Range factors were stratified as they were random factors

**Table S5.1.** Bacterial *amoA* Classification (inbred, hybrid, teosinte) model

| | Df | SumsOfSqs | MeanSqs | F.Model | R2 | Pr(>F) | |
|---|---|---|---|---|---|---|---|
| Type | 2 | 0.594 | 0.29722 | 0.86977 | 0.0056 | 0.661 | |
| Time | 2 | 1.691 | 0.84538 | 2.47386 | 0.01593 | 0.001 | *** |
| Row | 1 | 0.533 | 0.53302 | 1.55979 | 0.00502 | 0.102 | |
| Range | 1 | 0.413 | 0.41304 | 1.20869 | 0.00389 | 0.256 | |
| Block | 91 | 35.427 | 0.38931 | 1.13926 | 0.3337 | 0.007 | ** |
| Type:Time | 4 | 1.897 | 0.4742 | 1.38768 | 0.01787 | 0.028 | * |
| Residuals | 192 | 65.611 | 0.34172 | 0.618 | | | |
| Total | 293 | 106.166 | 1 | | | | |

**Table S5.2** Bacterial *amoA* Genotype Model

| | Df | SumsOfSqs | MeanSqs | F.Model | R2 | Pr(>F) | |
|---|---|---|---|---|---|---|---|
| Genotype | 26 | 9.403 | 0.36164 | 1.0811 | 0.08857 | 0.158 | |
| Time | 2 | 1.73 | 0.86514 | 2.5862 | 0.0163 | 0.001 | *** |
| Row | 1 | 0.533 | 0.53267 | 1.5923 | 0.00502 | 0.087 | . |
| Range | 1 | 0.396 | 0.39591 | 1.1835 | 0.00373 | 0.274 | |
| Block | 72 | 28.34 | 0.39362 | 1.1767 | 0.26694 | 0.007 | ** |
| Genotype:Time | 52 | 19.266 | 0.3705 | 1.1076 | 0.18147 | 0.042 | * |
| Residuals | 139 | 46.499 | 0.33452 | ng | | | |
| Total | 293 | 106.166 | 1 | | | | |

**Table S5.3** Bacterial *amoA* Nested Genotype/Type Model

| | Df | SumsOfSqs | MeanSqs | F.Model | R2 | Pr(>F) | |
|---|---|---|---|---|---|---|---|
| Classification | 2 | 0.594 | 0.29722 | 0.88849 | 0.0056 | 0.623 | |
| Time | 2 | 1.691 | 0.84538 | 2.52712 | 0.01593 | 0.001 | *** |
| Row | 1 | 0.533 | 0.53302 | 1.59338 | 0.00502 | 0.066 | . |
| Range | 1 | 0.413 | 0.41304 | 1.23471 | 0.00389 | 0.242 | |
| Block | 91 | 35.427 | 0.38931 | 1.16378 | 0.3337 | 0.001 | *** |
| Classification:Genotype | 5 | 1.743 | 0.34868 | 1.04231 | 0.01642 | 0.38 | |
| Classification:Time | 4 | 1.897 | 0.4742 | 1.41756 | 0.01787 | 0.033 | * |

| | | | | | | |
|---|---|---|---|---|---|---|
| Class:Genotype:Time | 48 | 17.369 | 0.36186 | 1.08171 | 0.1636 | 0.101 |
| Residuals | 139 | 46.499 | 0.33452 | 0.43798 | | |
| Total | 293 | 106.166 | 1 | | | |

**Table S5.4** Archeal *amoA* Classification Model

| | Df | SumsOfSqs | MeanSqs | F.Model | R2 | Pr(>F) | |
|---|---|---|---|---|---|---|---|
| Classification | 2 | 0.464 | 0.232 | 2.452 | 0.01165 | 0.008 | ** |
| Time | 2 | 6.806 | 3.403 | 35.955 | 0.17089 | 0.001 | *** |
| Row | 1 | 3.224 | 3.2242 | 34.066 | 0.08096 | 0.001 | *** |
| Range | 1 | 0.387 | 0.3867 | 4.086 | 0.00971 | 0.003 | ** |
| Block | 91 | 9.43 | 0.1036 | 1.095 | 0.23678 | 0.123 | |
| Classification:Time | 4 | 1.154 | 0.2886 | 3.049 | 0.02898 | 0.001 | *** |
| Residuals | 194 | 18.361 | 0.0946 | 0.46103 | | | |
| Total | 295 | 39.827 | 1 | | | | |

**Table S5.5** Archeal *amoA* Genotype Model

| | Df | SumsOfSqs | MeanSqs | F.Model | R2 | Pr(>F) | |
|---|---|---|---|---|---|---|---|
| Genotype | 26 | 3.346 | 0.1287 | 1.374 | 0.08401 | 0.007 | ** |
| Time | 2 | 6.798 | 3.3989 | 36.285 | 0.17068 | 0.001 | *** |
| Row | 1 | 2.708 | 2.7076 | 28.905 | 0.06799 | 0.001 | *** |
| Range | 1 | 0.406 | 0.4062 | 4.337 | 0.0102 | 0.002 | ** |
| Block | 72 | 7.812 | 0.1085 | 1.158 | 0.19614 | 0.027 | * |
| Genotype:Time | 52 | 5.55 | 0.1067 | 1.139 | 0.13935 | 0.075 | . |
| Residuals | 141 | 13.208 | 0.0937 | 0.33163 | | | |
| Total | 295 | 39.827 | 1 | | | | |

**Table S5.6** Archeal *amoA* nested Genotype/ Classification Model

| | Df | SumsOfSqs | MeanSqs | F.Model | R2 | Pr(>F) | |
|---|---|---|---|---|---|---|---|
| Classification | 2 | 0.464 | 0.232 | 2.477 | 0.01165 | 0.005 | ** |
| Time | 2 | 6.806 | 3.403 | 36.329 | 0.17089 | 0.001 | *** |
| Row | 1 | 3.224 | 3.2242 | 34.42 | 0.08096 | 0.001 | *** |
| Range | 1 | 0.387 | 0.3867 | 4.129 | 0.00971 | 0.003 | ** |
| Block | 91 | 9.43 | 0.1036 | 1.106 | 0.23678 | 0.101 | |
| Classification:Genotype | 5 | 0.758 | 0.1516 | 1.618 | 0.01903 | 0.024 | * |
| Classification:Time | 4 | 1.154 | 0.2886 | 3.081 | 0.02898 | 0.001 | *** |
| Class:Genotype:Time | 48 | 4.396 | 0.0916 | 0.978 | 0.11037 | 0.591 | |
| Residuals | 141 | 13.208 | 0.0937 | 0.33163 | | | |
| Total | 295 | 39.827 | 1 | | | | |

**Table S5.7** *nirK* Classification Model

| | Df | SumsOfSqs | MeanSqs | F.Model | R2 | Pr(>F) | |
|---|---|---|---|---|---|---|---|
| Classification | 2 | 1.11 | 0.555 | 1.43 | 0.00933 | 0.004 | ** |
| Time | 2 | 3.188 | 1.59391 | 4.1068 | 0.0268 | 0.001 | *** |
| Row | 1 | 0.688 | 0.68803 | 1.7727 | 0.00578 | 0.002 | ** |
| Range | 1 | 0.472 | 0.47213 | 1.2165 | 0.00397 | 0.09 | . |
| Block | 91 | 36.562 | 0.40178 | 1.0352 | 0.30733 | 0.042 | * |
| Classification:Time | 4 | 1.654 | 0.41339 | 1.0651 | 0.0139 | 0.21 | |
| Residuals | 194 | 75.295 | 0.38812 | 0.6329 | | | |
| Total | 295 | 118.968 | 1 | | | | |

**Table S5.8** *nirk* Genotype Model

| | Df | SumsOfSqs | MeanSqs | F.Model | R2 | Pr(>F) | |
|---|---|---|---|---|---|---|---|
| Genotype | 26 | 10.984 | 0.42248 | 1.0986 | 0.09233 | 0.003 | ** |
| Time | 2 | 3.142 | 1.57106 | 4.0854 | 0.02641 | 0.001 | *** |
| Row | 1 | 0.556 | 0.55561 | 1.4448 | 0.00467 | 0.011 | * |
| Range | 1 | 0.483 | 0.48293 | 1.2558 | 0.00406 | 0.064 | . |
| Block | 72 | 29.081 | 0.4039 | 1.0503 | 0.24444 | 0.019 | * |
| Genotype:Time | 52 | 20.5 | 0.39424 | 1.0252 | 0.17232 | 0.162 | |
| Residuals | 141 | 54.222 | 0.38455 | 0.45577 | | | |
| Total | 295 | 118.968 | 1 | | | | |

**Table S5.9** *nirK* Genotype/ Classification Model

| | Df | SumsOfSqs | MeanSqs | F.Model | R2 | Pr(>F) | |
|---|---|---|---|---|---|---|---|
| Classification | 2 | 1.11 | 0.555 | 1.4432 | 0.00933 | 0.001 | *** |
| Time | 2 | 3.188 | 1.59391 | 4.1448 | 0.0268 | 0.001 | *** |
| Row | 1 | 0.688 | 0.68803 | 1.7892 | 0.00578 | 0.001 | *** |
| Range | 1 | 0.472 | 0.47213 | 1.2277 | 0.00397 | 0.089 | . |
| Block | 91 | 36.562 | 0.40178 | 1.0448 | 0.30733 | 0.024 | * |
| Classification:Genotype | 5 | 2.226 | 0.44521 | 1.1577 | 0.01871 | 0.027 | * |
| Classification:Time | 4 | 1.654 | 0.41339 | 1.075 | 0.0139 | 0.159 | |
| Classification:Genotype:Time | 48 | 18.847 | 0.39264 | 1.021 | 0.15842 | 0.206 | |
| Residuals | 141 | 54.222 | 0.38455 | 0.45577 | | | |
| Total | 295 | 118.968 | 1 | | | | |

**Table S5.10** *nirS* Classification Model

| | Df | SumsOfSqs | MeanSqs | F.Model | R2 | Pr(>F) | |
|---|---|---|---|---|---|---|---|
| Classification | 2 | 1.303 | 0.65133 | 1.7675 | 0.01149 | 0.001 | *** |
| Time | 2 | 1.415 | 0.70762 | 1.9202 | 0.01248 | 0.001 | *** |
| Row | 1 | 1.224 | 1.22417 | 3.3219 | 0.0108 | 0.003 | ** |
| Range | 1 | 0.799 | 0.79936 | 2.1691 | 0.00705 | 0.001 | *** |
| Block | 91 | 35.535 | 0.3905 | 1.0597 | 0.31342 | 0.001 | *** |
| Classification:Time | 4 | 1.609 | 0.40222 | 1.0915 | 0.01419 | 0.185 | |
| Residuals | 194 | 71.492 | 0.36851 | 0.63056 | | | |
| Total | 295 | 113.377 | 1 | | | | |

**Table S5.11** *nirS* Genotype Model

| | Df | SumsOfSqs | MeanSqs | F.Model | R2 | Pr(>F) | |
|---|---|---|---|---|---|---|---|
| Genotype | 26 | 10.581 | 0.40697 | 1.08047 | 0.09333 | 0.031 | * |
| Time | 2 | 1.409 | 0.70462 | 1.8707 | 0.01243 | 0.001 | *** |
| Row | 1 | 1.015 | 1.01503 | 2.69481 | 0.00895 | 0.001 | *** |
| Range | 1 | 0.798 | 0.79755 | 2.11741 | 0.00703 | 0.001 | *** |
| Block | 72 | 28.471 | 0.39543 | 1.04984 | 0.25112 | 0.054 | . |
| Genotype:Time | 52 | 17.994 | 0.34603 | 0.91868 | 0.15871 | 0.998 | |
| Residuals | 141 | 53.109 | 0.37666 | 0.46843 | | | |
| Total | 295 | 113.377 | 1 | | | | |

**Table S5.12** *nirS* Nested Genotype/ Classification Model

| | Df | SumsOfSqs | MeanSqs | F.Model | R2 | Pr(>F) | |
|---|---|---|---|---|---|---|---|
| Classification | 1 | 0.34 | 0.3401 | 0.8887 | 0.00364 | 0.677 | |
| Time | 3 | 2.347 | 0.78227 | 2.04408 | 0.02512 | 0.001 | *** |
| Row | 1 | 0.98 | 0.97993 | 2.56059 | 0.01049 | 0.001 | *** |
| Range | 1 | 0.818 | 0.81839 | 2.13847 | 0.00876 | 0.002 | ** |
| Block | 80 | 31.504 | 0.39379 | 1.029 | 0.33723 | 0.171 | |
| Classification:Time | 2 | 0.748 | 0.37406 | 0.97742 | 0.00801 | 0.541 | |
| Class:Genotype:Time | 43 | 14.967 | 0.34808 | 0.90954 | 0.16022 | 0.999 | |
| Residuals | 109 | 41.714 | 0.3827 | 0.44653 | | | |
| Total | 240 | 93.418 | 1 | | | | |

**Table S5.13** *nosZ* Classification Model

| | Df | SumsOfSqs | MeanSqs | F.Model | R2 | Pr(>F) | |
|---|---|---|---|---|---|---|---|
| Classification | 2 | 1.288 | 0.64412 | 1.407 | 0.00935 | 0.001 | *** |
| Time | 2 | 1.857 | 0.92856 | 2.0283 | 0.01347 | 0.001 | *** |
| Row | 1 | 0.805 | 0.80456 | 1.7574 | 0.00584 | 0.001 | *** |
| Range | 1 | 0.581 | 0.58138 | 1.2699 | 0.00422 | 0.001 | *** |
| Block | 91 | 42.41 | 0.46604 | 1.018 | 0.30771 | 0.001 | *** |
| Classification:Time | 4 | 2.07 | 0.51758 | 1.1306 | 0.01502 | 0.013 | * |
| Residuals | 194 | 88.813 | 0.4578 | 0.64439 | | | |
| Total | 295 | 137.825 | 1 | | | | |

**Table S5.14** *nosZ* Genotype Model

| | Df | SumsOfSqs | MeanSqs | F.Model | R2 | Pr(>F) | |
|---|---|---|---|---|---|---|---|
| Genotype | 26 | 12.225 | 0.4702 | 1.0274 | 0.0887 | 0.011 | * |
| Time | 2 | 1.856 | 0.92801 | 2.0277 | 0.01347 | 0.001 | *** |
| Row | 1 | 0.786 | 0.78573 | 1.7168 | 0.0057 | 0.151 | |
| Range | 1 | 0.546 | 0.54594 | 1.1929 | 0.00396 | 0.057 | . |
| Block | 72 | 33.931 | 0.47126 | 1.0297 | 0.24619 | 0.034 | * |
| Genotype:Time | 52 | 23.951 | 0.4606 | 1.0064 | 0.17378 | 0.271 | |
| Residuals | 141 | 64.53 | 0.45766 | 0.4682 | | | |
| Total | 295 | 137.825 | 1 | | | | |

**Table S5.15** *nosZ* Genotype/ Classification Model

| | Df | SumsOfSqs | MeanSqs | F.Model | R2 | Pr(>F) | |
|---|---|---|---|---|---|---|---|
| Type | 2 | 1.288 | 0.64412 | 1.40742 | 0.00935 | 0.006 | ** |
| Time | 2 | 1.857 | 0.92856 | 2.02893 | 0.01347 | 0.001 | *** |
| Row | 1 | 0.805 | 0.80456 | 1.75799 | 0.00584 | 0.003 | ** |
| Range | 1 | 0.581 | 0.58138 | 1.27033 | 0.00422 | 0.008 | ** |
| Block | 91 | 42.41 | 0.46604 | 1.01831 | 0.30771 | 0.008 | ** |
| Classification:Genotype | 5 | 2.403 | 0.4805 | 1.04992 | 0.01743 | 0.193 | |
| Classification:Time | 4 | 2.07 | 0.51758 | 1.13093 | 0.01502 | 0.018 | * |
| Class:Genotype:Time | 48 | 21.881 | 0.45585 | 0.99605 | 0.15876 | 0.468 | |
| Residuals | 141 | 64.53 | 0.45766 | 0.4682 | | | |
| Total | 295 | 137.825 | 1 | | | | |

**Table S6.** Mixed effect models comparing qPCR of functional genes and genotype, plant classification, space, and time. Genotype and time of sampling were run as a fixed effect, while sampling block, range, and row were run as random factors. Wald tests performed on statistical models to calculate significance. Statistical models were run in ‘asreml-r’.

**Table S6.1** Bacterial *amoA* qPCR Classification Model

| Factor | Df | Sum of Sq | Wald | Pr(Chisq) | |
|---|---|---|---|---|---|
| (Intercept) | 1 | 167727 | 135.24 | 2.20E-16 | *** |
| Classification | 2 | 157 | 0.127 | 0.93856 | |
| Time | 2 | 10205 | 8.229 | 0.01634 | * |
| Classification:Time | 4 | 35000 | 28.221 | 1.13E-05 | *** |
| residual | | 1240 | | | |

**Table S6.5** Bacterial *amoA* qPCR Genotype Model

| Factor | Df | Sum of Sq | Wald | Pr(Chisq) | |
|---|---|---|---|---|---|
| (Intercept) | 1 | 159311 | 120.66 | 2.00E-16 | *** |
| Classification | 26 | 15361 | 11.634 | 0.99305 | |
| Time | 2 | 9378 | 7.103 | 0.02868 | * |
| Classification:Time | 52 | 83863 | 63.517 | 0.13144 | |
| residual | | 1320 | | | |

**Table S6.6** Arch *amoA* qPCR Classification Model

| Factor | Df | Sum of Sq | Wald | Pr(Chisq) | |
|---|---|---|---|---|---|
| (Intercept) | 1 | 468689956 | 37.761 | 8.00E-10 | *** |
| Classification | 2 | 35436473 | 2.855 | 0.2399 | |
| Time | 2 | 43443468 | 3.5 | 0.1738 | |
| Classification:Time | 4 | 76664810 | 6.177 | 0.1863 | |
| residual | | 12412117 | | | |

**Table S6.7** Arch *amoA* qPCR Genotype Model

| Factor | Df | Sum of Sq | Wald | Pr(Chisq) | |
|---|---|---|---|---|---|
| (Intercept) | 1 | 772154049 | 59.269 | 1.38E-14 | *** |
| Genotype | 26 | 305307733 | 23.435 | 0.6082 | |
| Time | 2 | 42938857 | 3.296 | 0.1924 | |
| **Genotype:**Time | 52 | 669965740 | 51.426 | 0.4964 | |
| residual | | 13027864 | | | |

**Table S6.8** *nirK* qPCR Classification Model

| Factor | Df | Sum of Sq | Wald | Pr(Chisq) | |
|---|---|---|---|---|---|
| (Intercept) | 1 | 3191339410 | 494.32 | 2.20E-16 | *** |
| Classification | 2 | 4872529 | 0.75 | 0.6856658 | |

| | | | | | |
|---|---|---|---|---|---|
| Time | 2 | 59666289 | 9.24 | 0.0098429 | ** |
| Classification:Time | 4 | 137359821 | 21.28 | 0.0002791 | *** |
| residual | | 6455991 | | | |

**Table S6.9** *nirK* qPCR Genotype Model

| Factor | Df | Sum of Sq | Wald | Pr(Chisq) | |
|---|---|---|---|---|---|
| (Intercept) | 1 | 3085673717 | 472.49 | 2.20E-16 | *** |
| Genotype | 26 | 113556098 | 17.39 | 0.896892 | |
| Time | 2 | 61635999 | 9.44 | 0.008925 | ** |
| Genotype:Time | 52 | 448758443 | 68.72 | 0.060059 | . |
| residual | | 6530707 | | | |

**Table S6.10** *nirS* qPCR Classification Model

| Factor | Df | Sum of Sq | Wald | Pr(Chisq) | |
|---|---|---|---|---|---|
| (Intercept) | 1 | 142927550 | 50.098 | 1.46E-12 | *** |
| Classification | 2 | 12327195 | 4.321 | 0.115275 | |
| Time | 2 | 91423589 | 32.045 | 1.10E-07 | *** |
| Classification:Time | 4 | 39142572 | 13.72 | 0.008244 | ** |
| residual | (MS) | 2852939 | | | |

**Table S6.11** *nirS* qPCR Genotype Model

| Factor | Df | Sum of Sq | Wald | Pr(Chisq) | |
|---|---|---|---|---|---|
| (Intercept) | 1 | 3085673717 | 472.49 | 2.20E-16 | *** |
| Genotype | 26 | 113556098 | 17.39 | 0.896892 | |
| Time | 2 | 61635999 | 9.44 | 0.008925 | ** |
| Genotype:Time | 52 | 448758443 | 68.72 | 0.060059 | . |
| residual | | 6530707 | | | |

**Table S6.12** *nosZ* qPCR Classification Model

| Factor | Df | Sum of Sq | Wald | Pr(Chisq) | |
|---|---|---|---|---|---|
| (Intercept) | 1 | 294866123 | 20.09 | 7.39E-06 | *** |
| Classification | 2 | 295357031 | 20.124 | 4.27E-05 | *** |
| Time | 2 | 1221569930 | 83.23 | 2.20E-16 | *** |
| Classification:Time | 4 | 719736962 | 49.038 | 5.73E-10 | *** |
| residual | | 14677047 | | | |

**Table S6.13** *nosZ* qPCR Genotype Model

| Factor | Df | Sum of Sq | Wald | Pr(Chisq) | |
|---|---|---|---|---|---|
| (Intercept) | 1 | 5127618121 | 308.503 | 2.00E-16 | *** |
| Genotype | 26 | 607829600 | 36.57 | 0.08167 | . |
| Time | 2 | 1227441338 | 73.849 | 2.00E-16 | *** |
| Genotype:Time | 52 | 1118209396 | 67.277 | 0.07547 | . |
| residual | | 16620956 | | | |

**Table S7.** Mixed effect models comparing potential nitrification and genotype, plant classification, space, and time. Genotype and growth stage were run as a fixed effect, while sampling block, sampling date, range, and row were run as random factors. Wald tests performed on statistical models to calculate significance. Statistical models were run in ‘asreml-r’.

**Table S7.1** Log of Nitrification Rate Per Genotype

| Terms | DF | Sum of Sq | Wald Statistic | Pr (Chisq) | Significant |
|---|---|---|---|---|---|
| (Intercept) | 1 | 25085.9 | 92498 | <2.2E-16 | *** |
| Genotype | 26 | 14.9 | 55 | 0.001 | ** |
| Growth Stage | 2 | 2.6 | 10 | 0.007 | ** |
| Genotype:Growth | 49 | 12.9 | 47 | 0.54 | |
| residual | | 0.3 | | | |

**Table S7.2** log of Nitrification Rate per Classification

| Terms | DF | Sum of Sq | Wald Statistic | Pr (Chisq) | Significant |
|---|---|---|---|---|---|
| (Intercept) | 1 | 7099.4 | 29956.9 | <2.2E-16 | *** |
| Classification | 2 | 4.7 | 19.8 | 5.11E-05 | *** |
| Growth Stage | 2 | 0.4 | 1.8 | 0.40 | |
| Classification:Growth | 4 | 3.1 | 13.2 | 0.01 | * |
| residual | | 0.2 | | | |

**Table S8.** Mixed effect models comparing potential denitrification and genotype, plant classification, space, and time. Genotype and growth stage were run as a fixed effect, while sampling block, range, sample date, and row were run as random factors. Wald tests performed on statistical models to calculate significance. Statistical models were run in 'asreml-r'.

**Table S8.1** Log of Overall Denitrification Enzyme Assay Genotype Model

| Terms | DF | Sum of Sq | Wald Statistic | Pr (Chisq) | Significant |
|---|---|---|---|---|---|
| (Intercept) | 1 | 22.01 | 16.539 | 4.77E-05 | *** |
| Genotype | 26 | 64.33 | 48.338 | 0.007 | ** |
| Growth Stage | 2 | 0.43 | 0.322 | 0.85 | |
| Genotype:Growth | 52 | 56.602 | 42.53 | 0.82 | |
| residual | | 1.33 | | | |

**Table S8.3** Log of Overall Denitrification Enzyme Assay Classification Model

| Terms | DF | Sum of Sq | Wald Statistic | Pr (Chisq) | Significant |
|---|---|---|---|---|---|
| (Intercept) | 1 | 11.27 | 8.84 | 2.9E-03 | ** |
| Classification | 2 | 22.02 | 17.28 | 2.7E-04 | *** |
| Growth Stage | 2 | 2.57 | 2.01 | 0.36 | |
| Classification:Growth | 4 | 4.89 | 3.84 | 0.43 | |
| residual | | 1.27 | | | |

**Table S8.4** Log of Incomplete Denitrification Enzyme Assay Genotype Model

| Terms | DF | Sum of Sq | Wald Statistic | Pr (Chisq) | Significant |
|---|---|---|---|---|---|
| (Intercept) | 1 | 1.71 | 0.96 | 0.32 | |
| Genotype | 26 | 56.32 | 31.55 | 0.24 | |
| Growth Stage | 2 | 6.58 | 3.689 | 0.16 | |
| Genotype:Growth | 52 | 119.30 | 66.24 | 0.09 | . |
| residual | | 1.79 | | | |

**Table S8.6** Log of Incomplete Denitrification Enzyme Assay Classification Model

| Terms | DF | Sum of Sq | Wald Statistic | Pr (Chisq) | Significant |
|---|---|---|---|---|---|
| (Intercept) | 1 | 3.49 | 1.91 | 0.166 | |
| Classification | 2 | 22.46 | 12.33 | 0.002 | *** |
| Growth Stage | 2 | 13.90 | 13.89 | 0.022 | * |
| Classification:Growth | 4 | 10.48 | 5.75 | 0.22 | |
| residual | | 1.82 | | | |

**Table S9.** Genotype effect calculations for potential nitrification, incomplete denitrification, and complete denitrification rates. Table includes genotype mean, variance, genotype effects which was calculated as the difference between the genotypes mean – the population mean, and the percent change which is the genotypic effects divided by the population mean. These metrics were adapted from Bernardo et al. *Breeding for Quantitative Traits in Plants.*

**Table S9.1** Potential Nitrification

| Genotypes | Classification | Mean | Variance | Genotype Effect | Percent Change |
|---|---|---|---|---|---|
| Ames21785 | Teosinte | 3223.202 | 1081182.57 | -691.15308 | -0.176568847 |
| Ames21786 | Teosinte | 2774.953 | 88433.44 | -1139.40144 | -0.291082838 |
| Ames21809 | Teosinte | 3133.105 | 746022.14 | -781.24917 | -0.199585692 |
| B73 | Inbred | 3932.59 | 2127444.5 | 18.23587 | 0.004658717 |
| B73/PHJ40 | Hybrid | 4063.868 | 2198144.3 | 149.51344 | 0.038196191 |
| B73/PHZ51 | Hybrid | 3307.112 | 1437775.21 | -607.24263 | -0.155132249 |
| CHECK1 | Hybrid | 3795.58 | 2885350.12 | -118.77462 | -0.030343348 |
| CHECK2 | Hybrid | 3889.966 | 1355353.96 | -24.38811 | -0.006230428 |
| LH1 | Inbred | 4412.906 | 1914200.4 | 498.55092 | 0.127364782 |
| LH123 | Inbred | 4243.162 | 675978.9 | 328.80741 | 0.084000415 |
| LH82 | Inbred | 4265.879 | 3109219.15 | 351.52435 | 0.089803911 |
| LH82/PH207 | Hybrid | 5856.937 | 28458139.93 | 1942.58205 | 0.496271353 |
| LH82/PHJ40 | Hybrid | 2983.589 | 3277029.92 | -930.76514 | -0.237782532 |
| Mo17 | Inbred | 4000.014 | 1853836.7 | 85.6593 | 0.021883378 |
| PH207 | Inbred | 4369.633 | 3803350.28 | 455.27798 | 0.116309845 |
| PHG35 | Inbred | 3714.55 | 3813926.52 | -199.8045 | -0.051044047 |
| PHG39 | Inbred | 3401.902 | 2167152.55 | -512.45228 | -0.130916162 |
| PHG39/PHZ51 | Hybrid | 3243.783 | 4451418.24 | -670.57131 | -0.171310824 |
| PHG47 | Inbred | 3825.386 | 3776140.43 | -88.96846 | -0.02272877 |
| PHG47/PHG84 | Hybrid | 4195.987 | 5260178.39 | 281.63267 | 0.071948687 |
| PHG47/PHJ40 | Hybrid | 3163.181 | 1001909.37 | -751.17393 | -0.191902373 |
| PHG84 | Inbred | 4927.571 | 3589665.08 | 1013.2166 | 0.258846401 |
| PHJ40 | Inbred | 4235.362 | 2139689.2 | 321.00759 | 0.082007795 |
| PHZ51 | Inbred | 3673.76 | 1545063.7 | -240.595 | -0.061464796 |
| PI566677 | Teosinte | 2987.754 | 1989648.34 | -926.60081 | -0.23671867 |
| PI658188 | Teosinte | 4287.181 | 26627889.28 | 372.82631 | 0.095245921 |
| PI658205 | Teosinte | 4870.039 | 26617163.44 | 955.68488 | 0.244148777 |

**Table S9.2** Overall Denitrification

| Genotypes | Classification | Mean | Variance | Genotype Effect | Percent Change |
|---|---|---|---|---|---|
| Ames21785 | Teosinte | 12.225411 | 779.658009 | -6.063614 | -0.33154387 |
| Ames21786 | Teosinte | 1.418217 | 7.923932 | -16.870808 | -0.92245529 |

| | | | | | |
|---|---|---|---|---|---|
| Ames21809 | Teosinte | 20.975168 | 4754.027636 | 2.686142 | 0.14687182 |
| B73 | Inbred | 26.411203 | 950.937289 | 8.122178 | 0.4441012 |
| B73/PHJ40 | Hybrid | 20.062967 | 1457.794312 | 1.773942 | 0.09699487 |
| B73/PHZ51 | Hybrid | 12.719734 | 120.740613 | -5.569291 | -0.30451545 |
| CHECK1 | Hybrid | 38.901194 | 4490.550549 | 20.612169 | 1.12702391 |
| CHECK2 | Hybrid | 11.884273 | 274.847049 | -6.404752 | -0.3501965 |
| LH1 | Inbred | 11.165771 | 161.450221 | -7.123254 | -0.38948243 |
| LH123 | Inbred | 12.949991 | 113.369208 | -5.339034 | -0.29192559 |
| LH82 | Inbred | 15.97309 | 642.964529 | -2.315935 | -0.12662977 |
| LH82/PH207 | Hybrid | 21.952365 | 250.834562 | 3.663339 | 0.20030261 |
| LH82/PHJ40 | Hybrid | 41.578123 | 1951.876885 | 23.289098 | 1.27339201 |
| Mo17 | Inbred | 25.755179 | 860.418624 | 7.466154 | 0.4082314 |
| PH207 | Inbred | 27.86796 | 1651.695357 | 9.578935 | 0.52375319 |
| PHG35 | Inbred | 23.894519 | 927.46544 | 5.605494 | 0.30649493 |
| PHG39 | Inbred | 8.609285 | 73.497137 | -9.67974 | -0.52926496 |
| PHG39/PHZ51 | Hybrid | 14.804714 | 201.503919 | -3.484311 | -0.19051378 |
| PHG47 | Inbred | 27.369599 | 2933.639208 | 9.080574 | 0.49650398 |
| PHG47/PHG84 | Hybrid | 25.585344 | 457.123644 | 7.296319 | 0.39894523 |
| PHG47/PHJ40 | Hybrid | 34.053087 | 1716.381409 | 15.764062 | 0.86194108 |
| PHG84 | Inbred | 24.404597 | 2230.25515 | 6.115572 | 0.3343848 |
| PHJ40 | Inbred | 15.48927 | 355.858064 | -2.799755 | -0.15308389 |
| PHZ51 | Inbred | 15.445163 | 208.991664 | -2.843862 | -0.15549553 |
| PI566677 | Teosinte | 3.194778 | 37.371736 | -15.094247 | -0.82531722 |
| PI658188 | Teosinte | 6.021103 | 68.286632 | -12.267923 | -0.67078056 |
| PI658205 | Teosinte | 1.354084 | 5.715189 | -16.934941 | -0.92596192 |

**Table S9.3** Incomplete Denitrification

| Genotypes | Classification | Mean | Variance | Genotype Effect | Percent Change |
|---|---|---|---|---|---|
| Ames21785 | Teosinte | 0.9733998 | 3.854093 | -7.98603912 | -0.891354826 |
| Ames21786 | Teosinte | 1.3051623 | 6.296865 | -7.6542766 | -0.85432544 |
| Ames21809 | Teosinte | 1.6059257 | 4.038354 | -7.35351324 | -0.820755999 |
| B73 | Inbred | 16.8188358 | 405.154126 | 7.85939689 | 0.877219763 |
| B73/PHJ40 | Hybrid | 7.4729687 | 48.611804 | -1.48647024 | -0.165911086 |
| B73/PHZ51 | Hybrid | 8.1838818 | 32.706687 | -0.7755571 | -0.086563133 |
| CHECK1 | Hybrid | 7.8783329 | 35.574311 | -1.08110598 | -0.120666705 |
| CHECK2 | Hybrid | 8.4449126 | 45.474519 | -0.51452636 | -0.057428413 |
| LH1 | Inbred | 8.6419164 | 55.18519 | -0.31752254 | -0.035440003 |
| LH123 | Inbred | 37.2180497 | 3699.631325 | 28.25861081 | 3.154060322 |
| LH82 | Inbred | 11.6609163 | 102.192998 | 2.70147736 | 0.301523051 |
| LH82/PH207 | Hybrid | 10.7900266 | 38.394728 | 1.83058766 | 0.204319454 |
| LH82/PHJ40 | Hybrid | 11.4109879 | 39.199536 | 2.45154898 | 0.273627512 |
| Mo17 | Inbred | 12.0360989 | 190.165106 | 3.07665997 | 0.343398733 |
| PH207 | Inbred | 8.5943683 | 34.988491 | -0.36507061 | -0.040747039 |

| | | | | | |
|---|---|---|---|---|---|
| PHG35 | Inbred | 8.2722852 | 49.910295 | -0.68715377 | -0.076696072 |
| PHG39 | Inbred | 9.8503228 | 80.708558 | 0.89088383 | 0.099435225 |
| PHG39/PHZ51 | Hybrid | 6.9874882 | 20.942325 | -1.97195076 | -0.220097573 |
| PHG47 | Inbred | 9.0023327 | 35.816546 | 0.04289375 | 0.004787548 |
| PHG47/PHG84 | Hybrid | 10.5552713 | 38.582969 | 1.59583234 | 0.178117441 |
| PHG47/PHJ40 | Hybrid | 9.0040367 | 74.346892 | 0.04459775 | 0.004977739 |
| PHG84 | Inbred | 9.5751494 | 39.744606 | 0.61571051 | 0.068721995 |
| PHJ40 | Inbred | 9.184837 | 40.428247 | 0.22539803 | 0.025157606 |
| PHZ51 | Inbred | 12.2326943 | 46.872466 | 3.27325542 | 0.365341563 |
| PI566677 | Teosinte | 2.7090858 | 8.303989 | -6.25035314 | -0.697627741 |
| PI658188 | Teosinte | 1.0254465 | 8.225691 | -7.93399238 | -0.885545674 |
| PI658205 | Teosinte | 5.9606611 | 106.052389 | -2.99877777 | -0.334705979 |

**Tables S10.** Permutational multivariate ANOVA model results at the genotypic level for 16S rRNA genes, and fungal ITS. Standard model was run on all amplicon sequence data. Bray-Curtis distance was used to calculate dissimilarity between microbiomes. 999 permutations were used in analysis. All factors in the model were run as fixed effect: ***Microbial Community Matrix* = Classification *x Sampling Time* + *Row x Range* + *Residuals.*** This model was used to all other nitrogen cycling genes compositional changes. Interactions between plant classification and time were incorporated to understand plant growth changes, while row and range interactions were used to understand spatial effects. Row and Range factors were stratified as they were random factors.

**Table 10.1** Classification (inbred, hybrid, teosinte) x Time Prokaryotic 16S rRNA Model

| Factor | Df | SumsOfSqs | MeanSqs | F.Model | R2 | Pr(>F) | |
|---|---|---|---|---|---|---|---|
| Classification | 2 | 1.8712 | 0.93559 | 12.4209 | 0.06447 | 0.001 | *** |
| Time | 2 | 2.0925 | 1.04623 | 13.8897 | 0.07209 | 0.001 | *** |
| Row | 1 | 1.0681 | 1.06807 | 14.1797 | 0.0368 | 0.001 | *** |
| Range | 1 | 0.3445 | 0.34454 | 4.5741 | 0.01187 | 0.001 | *** |
| Block | 91 | 8.3838 | 0.09213 | 1.2231 | 0.28884 | 0.001 | *** |
| Classification:Time | 4 | 1.1803 | 0.29507 | 3.9174 | 0.04066 | 0.001 | *** |
| Residuals | 187 | 14.0856 | 0.07532 | 0.48528 | | | |
| Total | 288 | 29.026 | 1 | | | | |

**Table 10.2** Genotype x Time Prokaryotic 16S rRNA Model (included all genotypes)

| Factor | Df | SumsOfSqs | MeanSqs | F.Model | R2 | Pr(>F) | |
|---|---|---|---|---|---|---|---|
| Genotype | 26 | 4.2653 | 0.16405 | 2.2087 | 0.14695 | 0.001 | *** |
| Time | 2 | 2.0737 | 1.03683 | 13.9596 | 0.07144 | 0.001 | *** |
| Row | 1 | 0.8968 | 0.89683 | 12.0748 | 0.0309 | 0.001 | *** |
| Range | 1 | 0.3204 | 0.32043 | 4.3142 | 0.01104 | 0.001 | *** |
| Block | 72 | 6.6467 | 0.09231 | 1.2429 | 0.22899 | 0.001 | *** |
| Genotype:Time | 52 | 4.8705 | 0.09366 | 1.2611 | 0.1678 | 0.001 | *** |
| Residuals | 134 | 9.9526 | 0.07427 | 0.34289 | | | |

| | | | |
|---|---|---|---|
| Total | 288 | 29.026 | 1 |

**Table 10.3** Nested Genotype-Type x Time Prokaryotic 16S rRNA Model

| Factor | Df | SumsOfSqs | MeanSqs | F.Model | R2 | Pr(>F) | |
|---|---|---|---|---|---|---|---|
| Type | 2 | 1.8712 | 0.93559 | 12.5966 | 0.06447 | 0.001 | *** |
| Time | 2 | 2.0925 | 1.04623 | 14.0862 | 0.07209 | 0.001 | *** |
| Row | 1 | 1.0681 | 1.06807 | 14.3803 | 0.0368 | 0.001 | *** |
| Range | 1 | 0.3445 | 0.34454 | 4.6388 | 0.01187 | 0.001 | *** |
| Block | 91 | 8.3838 | 0.09213 | 1.2404 | 0.28884 | 0.001 | *** |
| Type:Genotype | 5 | 0.4428 | 0.08855 | 1.1922 | 0.01525 | 0.069 | . |
| Type:Time | 4 | 1.1669 | 0.29174 | 3.9279 | 0.0402 | 0.001 | *** |
| Type:Genotype:Time | 48 | 3.7036 | 0.07716 | 1.0388 | 0.1276 | 0.164 | |
| Residuals | 134 | 9.9526 | 0.07427 | 0.34289 | | | |
| Total | 288 | 29.026 | 1 | | | | |

**Table 10.4** Classification (inbred, hybrid, teosinte) x Time Fungal ITS Model

| Factor | Df | SumsOfSqs | MeanSqs | F.Model | R2 | Pr(>F) | |
|---|---|---|---|---|---|---|---|
| Classification | 2 | 3.731 | 1.86552 | 6.0405 | 0.03629 | 0.001 | *** |
| Time | 2 | 2.267 | 1.13329 | 3.6696 | 0.02205 | 0.001 | *** |
| Row | 1 | 0.955 | 0.95467 | 3.0912 | 0.00929 | 0.001 | *** |
| Range | 1 | 0.527 | 0.52721 | 1.7071 | 0.00513 | 0.003 | ** |
| Block | 91 | 30.989 | 0.34054 | 1.1027 | 0.30143 | 0.001 | *** |
| Classification:Time | 4 | 4.423 | 1.10563 | 3.58 | 0.04302 | 0.001 | *** |
| Residuals | 194 | 59.914 | 0.30883 | 0.58279 | | | |
| Total | 295 | 102.805 | 1 | | | | |

**Table 10.5** Genotype x Time Fungal ITS Model (included all genotypes)

| Factor | Df | SumsOfSqs | MeanSqs | F.Model | R2 | Pr(>F) | |
|---|---|---|---|---|---|---|---|
| Genotype | 26 | 12.028 | 0.4626 | 1.5136 | 0.11699 | 0.001 | *** |
| Time | 2 | 2.246 | 1.12316 | 3.675 | 0.02185 | 0.001 | *** |
| Row | 1 | 0.995 | 0.99467 | 3.2546 | 0.00968 | 0.001 | *** |
| Range | 1 | 0.536 | 0.53647 | 1.7553 | 0.00522 | 0.007 | ** |
| Block | 72 | 24.303 | 0.33754 | 1.1044 | 0.2364 | 0.001 | *** |
| Genotype:Time | 52 | 19.604 | 0.377 | 1.2336 | 0.19069 | 0.001 | *** |
| Residuals | 141 | 43.093 | 0.30562 | 0.41917 | | | |
| Total | 295 | 102.805 | 1 | | | | |

**Table 10.6** Nested Genotype-Type x Time Fungal ITS Model

| | Df | SumsOfSqs | MeanSqs | F.Model | R2 | Pr(>F) | |
|---|---|---|---|---|---|---|---|
| Classification | 2 | 3.731 | 1.86552 | 6.1041 | 0.03629 | 0.001 | *** |

| | | | | | | | |
|---|---|---|---|---|---|---|---|
| Time | 2 | 2.267 | 1.13329 | 3.7082 | 0.02205 | 0.001 | *** |
| Row | 1 | 0.955 | 0.95467 | 3.1237 | 0.00929 | 0.001 | *** |
| Range | 1 | 0.527 | 0.52721 | 1.7251 | 0.00513 | 0.002 | ** |
| Block | 91 | 30.989 | 0.34054 | 1.1142 | 0.30143 | 0.001 | *** |
| Classification:Genotype | 5 | 1.64 | 0.32795 | 1.0731 | 0.01595 | 0.191 | |
| Classification:Time | 4 | 4.423 | 1.10563 | 3.6176 | 0.04302 | 0.001 | *** |
| Class:Genotype:Time | 48 | 15.181 | 0.31628 | 1.0349 | 0.14767 | 0.155 | |
| Residuals | 141 | 43.093 | 0.30562 | 0.41917 | | | |
| Total | 295 | 102.805 | 1 | | | | |

**Table S11.** The ASREML least-square means of significantly different soil chemistry analysis. Statistical significance was calculated using Wald's test $p<0.05$ and comparison was across plant classification.

| | Hybrid | Inbred | Teosinte |
|---|---|---|---|
| Buffer pH | 6.69±0.02 | 6.68±0.03 | 6.62±0.02 |
| Nitrate | 23.8± 4.75 | 27.89± 4.53 | 39.26± 3.20 |
| Phosphorus | 60.5±6.32 | 52.18±6.03 | 73.81±4.26 |
| Potassium | 184.9±36.56 | 189±34.86 | 348.04±24.65 |
| Sulfur | 8.1±0.65 | 8.09±0.62 | 10±0.44 |
| Manganese | 35.2±1.71 | 31.27±1.63 | 36.68±1.15 |
| Copper | 1.29±0.05 | 1.5±0.05 | 1.38±0.03 |
